# Epigenetic priming promotes acquisition of tyrosine kinase inhibitor resistance and oncogene amplification in human lung cancer

**DOI:** 10.1101/2025.01.26.634826

**Authors:** Rebecca M. Starble, Eric G. Sun, Rana Gbyli, Jonathan Radda, Jiuwei Lu, Tyler B. Jensen, Ning Sun, Nelli Khudaverdyan, Bomiao Hu, Mary Ann Melnick, Shuai Zhao, Nitin Roper, Gang Greg Wang, Jikui Song, Katerina Politi, Siyuan Wang, Andrew Z. Xiao

## Abstract

In mammalian cells, gene copy number is tightly controlled to maintain gene expression and genome stability. However, a common molecular feature across cancer types is oncogene amplification, which promotes cancer progression by drastically increasing the copy number and expression of tumor-promoting genes. For example, in tyrosine kinase inhibitor (TKI)-resistant lung adenocarcinoma (LUAD), oncogene amplification occurs in over 40% of patients’ tumors. Despite the prevalence of oncogene amplification in TKI-resistant tumors, the mechanisms facilitating oncogene amplification are not fully understood. Here, we find that LUADs exhibit a unique chromatin signature demarcated by strong CTCF and cohesin deposition in drug-naïve tumors, which correlates with the boundaries of oncogene amplicons in TKI-resistant LUAD cells. We identified a global chromatin priming effect during the acquisition of TKI resistance, marked by a dynamic increase of H3K27Ac, cohesin loading, and inter-TAD interactions, which occurs before the onset of oncogene amplification. Furthermore, we have found that the METTL7A protein, which was previously reported to localize to the endoplasmic reticulum and inner nuclear membrane, has a novel chromatin regulatory function by binding to amplified loci and regulating cohesin recruitment and inter-TAD interactions.

Surprisingly, we discovered that METTL7A remodels the chromatin landscape prior to large-scale copy number gains. Furthermore, while METTL7A depletion has little effect on the chromatin structure and proliferation of drug-naïve cells, METTL7A depletion prevents the formation and maintenance of TKI resistant-clones, highlighting the specific role of METTL7A as cells are becoming resistant. In summary, we discovered an unexpected mechanism required for the acquisition of TKI resistance regulated by a largely uncharacterized factor, METTL7A. This discovery sheds light into the maintenance of oncogene copy number and paves the way to the development of new therapeutics for preventing TKI resistance in LUAD.

## Main

Acquired resistance to tyrosine kinase inhibitors (TKIs) such as osimertinib, a third generation TKI, is a major clinical challenge in the treatment of EGFR-mutant non-small cell lung cancer (NSCLC). One common mechanism of TKI resistance is oncogene amplification^1–5^, which occurs in over 40% of TKI-resistant lung adenocarcinoma (LUAD) tumors harboring EGFR mutations^6^. Oncogene amplification can occur both intrachromosomally and through the form of extrachromosomal DNA (ecDNA)^7–12^. Despite the prevalence of oncogene amplification in TKI-resistant tumors, it is still not fully understood how chromatin architecture changes throughout the acquisition of resistance to facilitate oncogene amplification. Furthermore, while previous studies have demonstrated that TKI-resistant tumors emerge from a quiescent, drug-tolerant persister (DTP) stage characterized by a unique and reversible epigenetic state^13^, the mechanisms that govern the exit from the DTP stage and entry into a proliferative, drug-resistant stage remain elusive. Here, we identify a novel chromatin regulator, METTL7A, that is specifically upregulated as LUAD cells exit the DTP stage and acquire resistance to osimertinib, during which METTL7A primes the epigenetic landscape of future amplicons through the recruitment of the cohesin complex.

## TKI-resistant LUAD tumors exhibit diverse forms of oncogene amplicons

To understand the genomic and structural changes that occur when LUAD tumors acquire resistance to osimertinib, we first performed whole genome sequencing (WGS) in the paired parental and osimertinib-resistant (OR) EGFR-mutant LUAD cell lines PC9/PC9-OR, HCC827/HCC827-OR, and H1975/H1975-OR (**Extended Data Fig. 1a**). Through AmpliconArchitect (AA) analysis^14^, we observed gene amplification in both parental and resistant cells, which was exacerbated in resistant cells. In both PC9 and HCC827 cell lines, we observed increased gene amplification in the resistant cells compared to their parental counterparts (**Fig. 1a, b**), which is consistent with oncogene amplification being a common feature of osimertinib resistance in patient tumors^6^. Consistent with previous work^15^, *RAF1* was one of the most highly amplified oncogenes in PC9-OR cells, which was predicted by AA to exhibit an extrachromosomal DNA (ecDNA) structure. In HCC827 cells, we identified that *EGFR* was highly amplified in both parental and osimertinib-resistant cells, with increased copy numbers in HCC827-OR cells. Furthermore, we found that HCC827-OR cells exhibited *MET* amplification, which is consistent with previous reports that have identified *MET* amplification as a mechanism of gefitinib resistance in the HCC827 cell line^4^ and osimertinib resistance in patient tumors^6^. In H1975 cells, we observed *PVT1-MYC* amplification in both parental and resistant cells, and this oncogene exhibited a breakage-fusion-bridge (BFB) signature. In addition, we examined the complexity of amplicons in these paired cell lines and observed that osimertinib-resistant cells had a consistent trend toward an increase in amplicon complexity score, a metric that accounts for the size and number of amplicons^16^ (**Extended Data Fig. 1b**). We validated these WGS results through DNA fluorescence in situ hybridization (FISH) in interphase nuclei using probes targeting amplified oncogenes and non-amplified chromosomal control loci (**Fig. 1c, Extended Data Fig. 1c, d**). Metaphase FISH revealed that most amplified oncogenes adopt the form of homogenously-staining regions (HSRs), a structure that can arise from ecDNA reintegration into chromosomes (**Extended Data Fig. 1e**), compared to extrachromosomal amplification, which has been observed in cell lines such as COLO320-DM (**Extended Data Fig. 1f**). To determine whether increased amplification correlates with increased transcription, we performed RNA sequencing (RNA-seq) and correlated these results with our AA analysis. Indeed, the amplified oncogenes were some of the most highly expressed genes in the transcriptome, including *RAF1* in PC9-OR and *MET* in HCC827-OR (**Fig. 1d, Supplementary Table 1**).

**Fig. 1.**
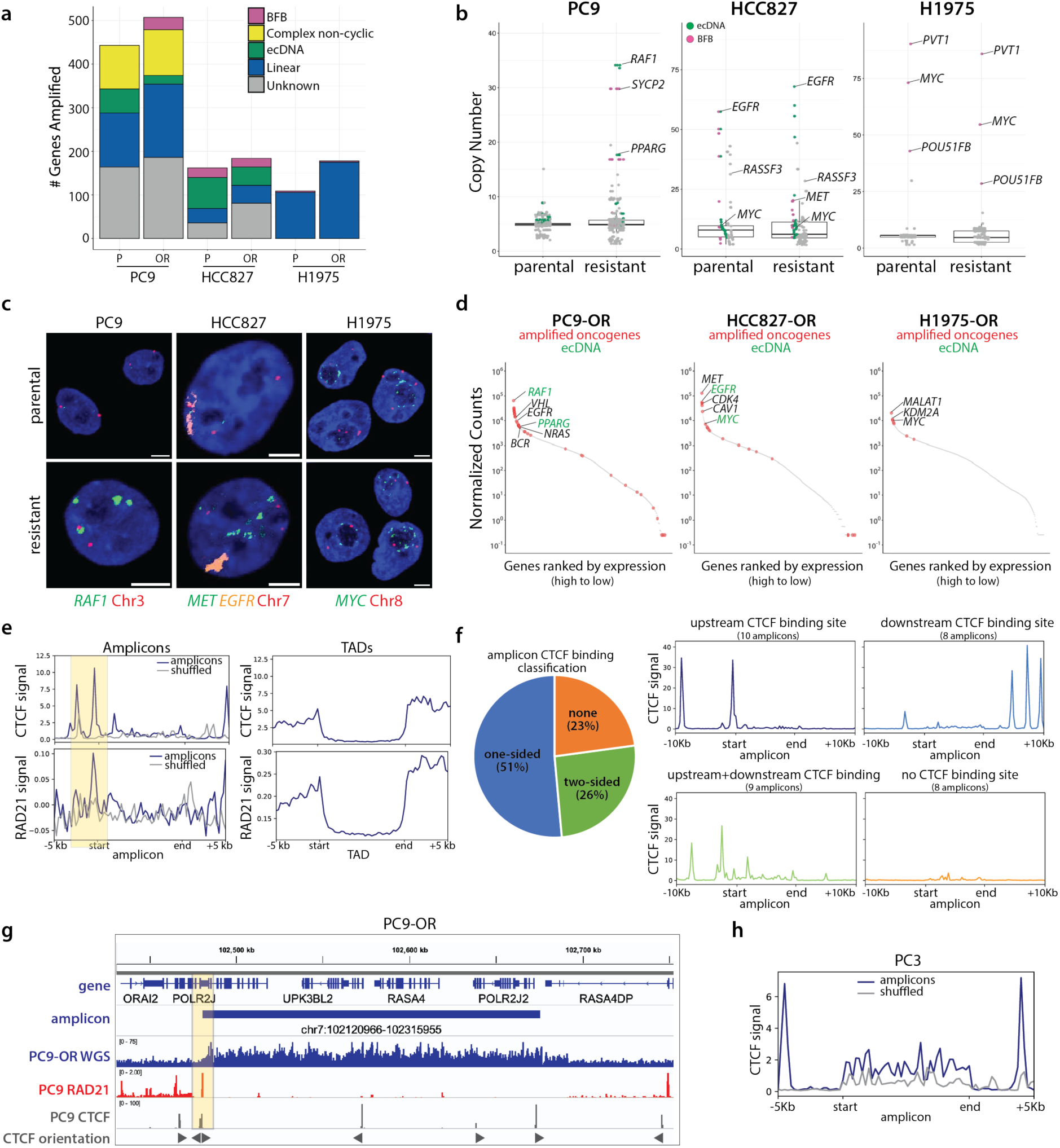
**Osimertinib-resistant EGFR-mutant LUADs exhibit oncogene amplicons demarcated by CTCF binding**. **a,** Whole-genome sequencing (WGS) and AmpliconArchitect (AA) analysis of parental and osimertinib-resistant cells shows the breakdown of amplicon structures across cell lines. P: parental, OR: osimertinib-resistant. **b**, Scatterplot of genes amplified across cell lines reveals an increase in amplicon copy number in PC9-OR and HCC827-OR cells compared to their parental counterparts. Pink and green dots denote genes predicted to be amplified via breakage-fusion-bridge (BFB) cycles and as extrachromosomal DNA (ecDNA) respectively based on AA analysis. **c,** Representative FISH images of amplified oncogenes and unamplified, control chromosomal loci. Scale bars: 5 µm. **d,** Ranked RNA expression plots in osimertinib-resistant cell lines. Red dots: amplified oncogenes; green dots: ecDNA based on WGS/AA analysis. **e,** Analysis of average CTCF and RAD21 signal at the boundaries of amplicons (left) compared to average signal at TAD boundaries (right) in PC9 cells. CTCF and RAD21 signal is increased at amplicon boundaries. Amplicon boundaries are determined by AA/WGS analysis in PC9-OR cells and TAD boundaries are determined by PC9 Hi-C. Because amplicons range in size from approximately 15 kb to 10 Mb, CTCF signal was plotted over scaled amplicons. Intersection between CTCF peaks and amplicons: p < 7.5 x 10^-15^, Fisher’s exact test. **f,** Left: amplicon CTCF motif classification based on the presence of a CTCF motif within 10kb of the start and/or end of the amplified interval. CTCF ChIP data is from PC9 parental cells; amplicons are based on AA from PC9-OR cells. Right: PC9 CTCF signal plotted over each of the PC9-OR amplicons categorized by the presence of CTCF motifs. **g,** Example IGV tracks showing CTCF (light grey) and RAD21 (red) enrichment over amplicon (dark blue) boundaries. CTCF orientation is indicated by grey triangles. **h,** CTCF signal plotted over amplicons (or amplicon coordinates shuffled over the genome, grey) in PC3 cell line.

We next examined the prevalence of oncogene amplification in osimertinib-resistant EGFR-mutant LUADs from patients (NCT02759835) from whole exome sequencing (WES) and RNA-seq data. We examined RNA expression in tumors which progressed upon osimertinib and performed copy number analysis on osimertinib-resistant tumors compared to pre-osimertinib-treated tumors. Consistent with previous results^17^, we identified 4 tumors that developed MET amplification and 5 that developed EGFR amplification in patient samples. We correlated this data with RNA expression and found that similar to our analyses in cell lines, these amplified genes were often the most highly expressed genes in the transcriptome (**Extended Data Fig. 2**).

To further understand the structure of the oncogene amplicons in osimertinib-resistant patient tumors, we performed WGS and AA analysis on patient-derived xenograft (PDX) models from a subset of osimertinib-treated patients from NCT02759835^18^ (**Supplementary Table 2**). Of the 9 TKI-treated PDXs we sequenced, 4/9 tumors exhibited high levels of oncogene amplification (>50 genes amplified), 3/9 tumors exhibited low levels of oncogene amplification (<50 genes amplified), and 2/9 tumors exhibited no gene amplification as detected by AA (**Extended Data Fig. 3a, b**). Across the PDXs, the most frequently amplified oncogenes were *EGFR, MDM2, KRAS,* and *MET* (**Extended Data Fig. 3c**). Of the 4 resistant tumors with high oncogene amplification, diverse amplicon structures were detected by AA, such as linear amplifications, complex rearrangements, ecDNA, and breakage-fusion bridge structures. These amplified oncogenes correlated strongly with high transcription based on RNA-seq (**Extended Data Fig. 3d**). Furthermore, these tumors exhibited high amplicon complexity scores, suggesting that these amplicons may have undergone successive cycles of amplification, recombination, and integration during tumor evolution (**Extended Data Fig. 3e**).

Together, using *in vitro*, PDX, and clinical trial data, our results indicate that oncogene amplification is highly prevalent in osimertinib-resistant EGFR-mutant LUAD, and these amplicons occur in the form of diverse chromosomal and extrachromosomal structures.

## A unique chromatin landscape signature in drug-naïve EGFR-mutant LUAD correlates with oncogene amplicon boundaries in TKI-resistant LUAD and other tumor types

While we observed a diverse range of amplicon structures across osimertinib-resistant cell lines and PDX LUAD tumors, we hypothesized that there could be a unique chromatin signature that permits the amplification of oncogene amplicons regardless of each amplicon’s unique oncogene locus or structure. Alterations in chromatin topology have been well documented in cancer^19–23^, and these alterations can be caused by changes in the binding of CCCTC-Binding factor (CTCF), which regulates chromatin structure in part due to its role in demarcating topologically associated domains (TADs), also known as contact domains. For example, CTCF was recently shown to affect the copy number and rearrangement of the *KMT2A* gene amplification in leukemia^24^. Thus, we sought to determine whether amplicon structure is associated with CTCF. We analyzed publicly available CTCF chromatin immunoprecipitation sequencing (ChIP-seq, ENCSR243INX) and Hi-C data (ENCSR859DRK) from the ENCODE Project^25,26^ performed in parental PC9 cells that were never exposed to TKIs. We correlated CTCF binding in PC9 cells with our PC9-OR WGS data and observed that CTCF was enriched at the boundaries of amplicons determined by AA compared to randomly shuffled, non-amplicon coordinates over the genome and this correlation is statistically significant (p < 7.5 x 10^-15^; **Fig. 1e, left**). This finding is particularly surprising because this CTCF enrichment occurred in drug-naïve PC9 cells that lack detectable levels of oncogene amplification (**Fig. 1b-d**). Thus, we deemed these loci “future amplicons” based on the observation that CTCF is enriched over the boundaries of these loci in cells which have not yet gained amplicons at these sites. To determine the strength of the CTCF signal at these “future amplicon” boundaries, we compared these data to CTCF signal at TAD boundaries, annotated based on the PC9 Hi-C data from ENCODE. Indeed, CTCF signal over amplicon boundaries was even more pronounced than at TAD boundaries (**Fig. 1e, right**). Interestingly, we observed that CTCF was frequently enriched on one side of the amplicon boundary, which is a similar feature to super-stripes, which facilitate long-range interactions across TAD boundaries^27^. This one-sided CTCF enrichment contrasts the enriched CTCF signal at both ends of TAD boundaries (**Fig. 1e**). To determine whether CTCF enrichment occurs at loop anchor sites, we performed CUT&RUN for the cohesin component RAD21 in drug-naïve PC9 cells and correlated this data with the CTCF data. We observed a similar enrichment of RAD21 over the “future amplicon” boundaries, with a similarly strong bias toward enrichment over one end of the amplicon (**Fig. 1e, bottom**).

To further study the trend of one-sided CTCF enrichment, we classified amplicons into the following 4 categories based on the presence of at least one CTCF binding site within 10kb upstream or downstream of the amplicon edges: amplicons exhibiting an upstream CTCF binding site (10 amplicons), downstream CTCF binding site (8 amplicons), CTCF binding sites at both boundaries (9 amplicons), or no CTCF binding sites (8 amplicons) (**Fig. 1f, left**). We plotted CTCF signal over amplicons within each category and observed a significant overlap between the presence of a CTCF binding site within 10kb of the amplicon boundary (Fisher’s Exact Test, p < 8.5 x 10^-12^), with 18/35 amplicons exhibiting this one-sided enrichment (**Fig. 1f, right**). Even for the amplicons characterized by “two-sided” CTCF binding (9/35), CTCF signal was still much stronger at one amplicon boundary compared to the other (**Fig. 1f, right**). When we examined individual amplicons more closely, this similarity to architectural super-stripes became even more apparent. For example, PC9-OR cells exhibited an amplicon on chromosome 7 containing the genes *UPK3BL2, RASA4,* and *POLR2J2*. The upstream boundary of this amplicon contained a strong CTCF binding site, oriented towards the amplicon, which overlapped with RAD21 deposition (**Fig. 1g**). This could indicate the presence of a pre-existing loop anchor in drug-naïve cells, which may facilitate the formation of this amplicon upon osimertinib treatment.

To further study this trend, we correlated CTCF binding in PC9 cells with the amplified intervals in the PDX tumors derived from osimertinib-treated tumors previously described. In the two PDX tumors (LAT001 and TMN0123) with the highest levels of oncogene amplification based on WGS and AA (**Extended Data Fig. 3a**), we observed that CTCF signal was similarly enriched over the amplicon boundaries, with a biased enrichment near one side of the amplicon (**Extended Data Fig. 3f, g**). This further indicates that the loci that become amplified during the acquisition of osimertinib resistance may have a predetermined higher-order chromatin architecture that already exists within the drug-naïve tumor. Together, these data suggest that prior to oncogene amplification, a unique chromatin landscape signature may pre-exist at the “future amplicon” boundaries, which may be regulated via cohesin-mediated loop extrusion.

To determine whether this observation more broadly occurs in different tumor types, we re- analyzed WGS data from PC3^7^ prostate cancer cells and vemurafenib (BRAF inhibitor)- and selumetinib (MEK inhibitor)-resistant, *BRAF^V^*^600^*^E^*-mutant M249 melanoma cells^28^. We correlated these data with CTCF ChIP-seq performed in PC3 (ENCSR359LOD) and neural crest cells (ENCSR218MVT), respectively, from the ENCODE Project^25,26^. We decided to analyze data from neural crest cells because they are the progenitor cells for melanocytes, which eventually give rise to melanoma. Thus, we reasoned that CTCF deposition in neural crest cells may more closely resemble melanocytes than other tissue types. AA analysis of PC3 cells revealed 35 amplified intervals, with an average copy number of 10. The most prominent oncogene amplified in this cell line was *PVT1-MYC*, which has previously been shown to exist as ecDNA. As previously reported^28^, we observed that vemurafenib- and selumetinib-resistant M249 cells contained only one amplicon, which contained the *BRAF* locus. Similar to the trend in LUAD tumors, we observed enriched CTCF signal at amplicon boundaries in both PC3 (**Fig. 1h**) and M249 cell lines (**Extended Data Fig. 3h**). While vemurafenib- and selumetinib-resistant M249 cells only contained one amplicon (*BRAF*), we were particularly intrigued by the observation that CTCF was enriched at this locus in neural crest cells, which do not harbor gene amplification.

Together, these intriguing observations suggest that non-amplified, drug-naïve tumor cells may already have a predefined chromatin structure that is permissive to the future amplification of these loci, a phenomenon that may occur in multiple types of oncogene-amplified cancers.

## Dynamic changes in chromatin architecture precede oncogene amplification during the acquisition of resistance

Previous studies demonstrated that TKI-resistant tumors emerge from a quiescent, drug-tolerant persister (DTP) stage characterized by a unique and reversible epigenetic state^13^. Although histone demethylation has been implicated as an acute response to TKIs^29^, the mechanisms that govern the exit from the DTP stage and entry into a proliferative, drug-resistant stage, which is a critical step to form resistant clones, remain elusive (**Fig. 2a**). Furthermore, the time point at which oncogene amplification occurs during the acquisition of resistance in EGFR-mutant LUAD is also unknown. To address these questions, we first leveraged a time course experiment in which we treated PC9 parental cells with escalating doses of osimertinib (100 nM to 1 μM) over the course of 3 months and probed for oncogene amplification. We first examined global copy number by plotting the normalized sequencing read coverage over each PC9-OR amplicon locus at 3 time points: PC9 cells treated with osimertinib for 0, 6-8, or 12 weeks. After 6-8 weeks of osimertinib treatment, PC9 cells had started to enter an exponential growth stage, and after 12 weeks, cells were stably resistant to osimertinib. We started to observe a global increase in copy number after approximately 12 weeks of osimertinib treatment (**Extended Data Fig. 4a**). We next probed for amplification of the *RAF1* locus at these same points at the single-cell level by DNA FISH. Consistent with the bulk sequencing data, we observed that global *RAF1* amplification occurred after 12 weeks of osimertinib treatment (**Fig. 2b, c**). At the “early resistance” 6–8-week stage, we observed a modest yet significant increase in *RAF1* copy number, suggesting that this is the time point immediately preceding global increases in gene copy number.

**Fig. 2.**
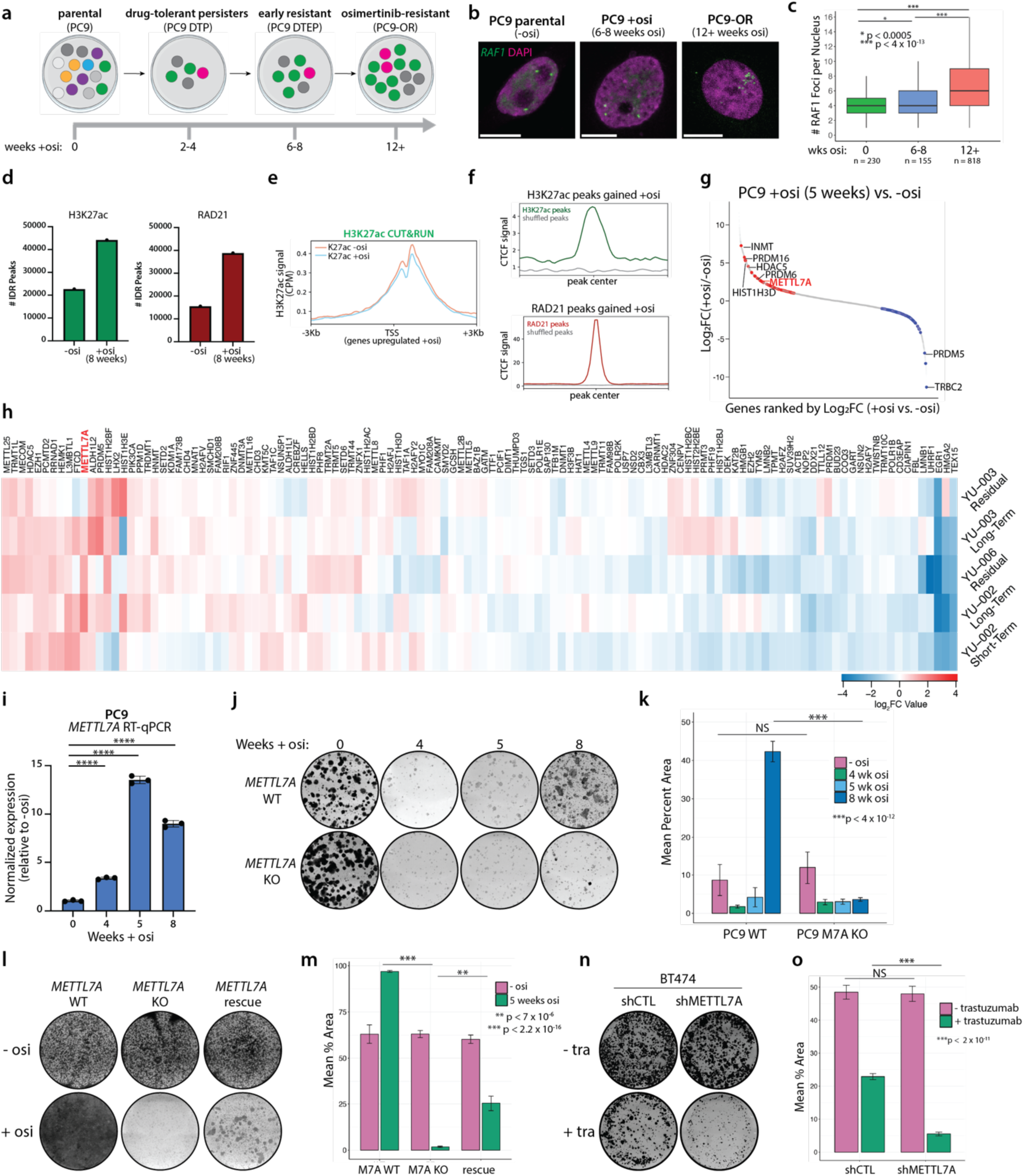
METTL7A promotes the acquisition of osimertinib-resistant LUAD. **a,** Schematic of PC9 cell line model of acquired osimertinib resistance. Cells transition through a quiescent, drug-tolerant persister (DTP) state before entering a proliferative resistant state. **b,** DNA FISH for the *RAF1* locus (green) in PC9 cells treated for 0, 8, or 12+ weeks with osimertinib. **c,** Quantification of Fig. 2b. Significance determined by unpaired t-test. N = number of nuclei imaged per sample from two biological replicates. **d,** Total number of significant H3K27ac peaks (left) and RAD21 peaks (right) in PC9 cells treated without osimertinib or for 8 weeks with osimertinib. Plotted are the consensus peaks based on IDR analysis between two biological replicates. **e,** H3K27ac CUT&RUN signal in PC9 cells treated without osimertinib or with osimertinib for 6-8 weeks, centered on genes significantly upregulated upon osimertinib treatment. H3K27ac signal is normalized by CPM and H2A is subtracted. Gain of H3K27ac signal in 8-week osimertinib-treated cells is not correlated with differentially expressed genes. **f,** CTCF signal is enriched at H3K27ac and RAD21 peaks gained upon osimertinib treatment compared to peaks randomly shuffled over the genome. **g,** Ranked RNA expression plot showing the differentially expressed (P_adj_ < 0.001) putative and known chromatin and epigenetic factors that are upregulated (red, log_2_FC > 1) or downregulated (blue, log_2_FC < 1) in PC9 cells after 5 weeks of osimertinib treatment. **h,** Heatmap of differentially expressed epigenetic factors in PDX models treated with osimertinib compared to vehicle. *METTL7A* (red) is one of the top upregulated genes across PDX samples treated with osimertinib. **i,** RT-qPCR validation of RNA- seq shows significant upregulation of *METTL7A* during the acquisition of osimertinib resistance in PC9 cells. *P* < 0.0001, calculated via an unpaired t-test. Standard deviation is based on three technical replicates. **j,** Long-term colony formation assay in PC9 WT and *METTL7A* KO cells stained with crystal violet at the indicated time points during chronic (1 µM) osimertinib treatment. **k,** Quantification of 2j. Error bars represent standard deviation between three biological replicates. **l,** Colony formation assay of PC9 WT, METTL7A KO, and METTL7A rescue cells treated with increased doses of osimertinib (0.1 to 1 µM) for 5 weeks. METTL7A (M7A) rescue cells were derived by overexpressing METTL7A in the KO background. **m**, Quantification of colony formation assay. Error bars represent standard deviation between biological triplicates. Significance determined by unpaired t-test. **n,** Crystal violet assay performed in shCTL or shMETTL7A BT474 breast cancer cells treated with or without trastuzumab. **o,** Quantification of (n). Error bars represent standard deviation between biological triplicates. Significance determined by unpaired t-test.

We next aimed to elucidate how chromatin architecture changes during the acquisition of osimertinib resistance prior to oncogene amplification. Since histone acetylation changes are associated with the DTP stage^13,29^, we first performed CUT&RUN for H3K27ac, a marker for enhancers poised for activation. In PC9 cells treated with osimertinib for 6-8 weeks, the time point before global oncogene amplification occurs, we observed a global increase in H3K27ac compared to drug-naïve PC9 cells (**Fig. 2d, left**). Interestingly, even though this time point occurs before the onset of oncogene amplification, we observed a significant intersection between the H3K27ac peaks gained with osimertinib treatment after 6-8 weeks and the oncogene amplicons in PC9-OR cells (p < 2 x 10^-10^). We next wanted to know whether the gain of H3K27ac peaks corresponded to changes in gene expression. Interestingly, although H3K27ac is frequently associated with transcriptional upregulation, further analysis revealed that the gained H3K27ac peaks were not correlated with concurrent gene expression level changes (**Fig. 2e, Extended Data Fig. 4b**). Instead, the gained H3K27ac peaks were associated with CTCF deposition from PC9 parental cells (**Fig. 2f**). Based on this observation, we hypothesized that chromatin undergoes changes in cohesin-mediated chromatin looping during the acquisition of resistance. Thus, we performed CUT&RUN for the core cohesin component RAD21. Similar to our H3K27ac data, we observed a global increase in RAD21 deposition in PC9 cells treated with osimertinib for 6-8 weeks (**Fig. 2d, right**), which was similarly correlated with CTCF signal (**Fig. 2f**). These data suggest that chromatin undergoes dynamic changes that precede massive oncogene amplification as tumors acquire resistance to osimertinib. We hypothesized that these changes occur at CTCF sites in drug-naïve cells, indicative of a chromatin priming event.

We next sought to determine the functional effects of oncogene amplification during the acquisition of resistance. To address this, we employed an shRNA-mediated knockdown strategy. We infected PC9 and HCC827 parental cells with shRNAs targeting either *RAF1* or *MET,* respectively, or a non-targeting control shRNA and treated cells with osimertinib over the course of approximately one month (**Extended Data Fig. 4c**). We found that both *RAF1* and *MET* knockdown resulted in significantly fewer osimertinib-resistant clones in PC9 and HCC827 cells, respectively, compared to control cells (p < 2.2 x 10^-16^ and p < 2 x 10^-7^, respectively; **Extended Data Fig. 4d, e**). This data further emphasizes the functional role of oncogene amplification during the acquisition of osimertinib resistance.

## METTL7A promotes the acquisition of osimertinib resistance

We next sought to identify novel chromatin regulators that facilitate chromatin reorganization prior to oncogene amplification. We were particularly interested in factors upregulated after 5 weeks of osimertinib treatment because this is a critical time point corresponding to the exit from the DTP stage. Thus, we performed RNA-seq in PC9 cells at multiple time points during the acquisition of osimertinib resistance. Given the strong change in chromatin landscape during the acquisition of resistance, we specifically profiled known and putative chromatin binding and or epigenetic factors (see **Methods** for details) for differential expression between untreated and osimertinib-treated cells at multiple time points during the acquisition of resistance (0, 4, 5, or 8 weeks of treatment). At 5-weeks post treatment, we identified 43 factors that were significantly upregulated (p_adj_ < 0.001, log_2_FC > 1) in osimertinib-treated cells compared to parental PC9 cells (**Fig. 2g**). Similarly, we identified 39 putative and known epigenetic factors that were significantly upregulated at both the 4-week and 8-week time points (**Extended Data Fig. 5a**).

To hone in on the factors most essential for the acquisition of resistance in a physiologically-relevant context, we analyzed RNA-seq in matched vehicle- and osimertinib-treated PDX tumors, consisting of PDXs treated with osimertinib for approximately 3 days (short-term), multiple months (long-term), and residual tumors that have persisted after osimertinib treatment^46^. We were particularly interested in differentially upregulated, putative epigenetic factors in both the cell line and PDX datasets. One of the top shared, differentially upregulated genes that emerged was *METTL7A*, which encodes Methyltransferase-like protein 7A (METTL7A). *METTL7A* was significantly upregulated during the acquisition of resistance in PC9 cells treated at all 3 time points (4, 5, and 8 weeks post-osimertinib treatment) and in 4/5 osimertinib-treated PDX tumors (**Fig. 2h**). We validated this result with qRT-PCR in PC9 cells treated with osimertinib, which similarly showed that *METTL7A* expression peaked after 5 weeks of osimertinib treatment (**Fig. 2i**). We additionally validated that this upregulation occurs in HCC827 cells (**Extended Data Fig. 5b**).

We next sought to address whether *METTL7A* upregulation arises from a preexisting, *METTL7A*-high population or is induced upon osimertinib treatment. We analyzed published scRNA-seq data performed in untreated and osimertinib-treated, drug-tolerant residual cells from an EGFR-mutant lung adenocarcinoma PDX, YU-006^46^. We observed that *METTL7A* expression was enriched in cluster 8, which is predominantly composed of osimertinib-treated, drug-tolerant cells. Unlike untreated cells (blue) in clusters 2 and 8, *METTL7A* expression was specific to the osimertinib-treated, drug-tolerant residual cells in cluster 8 (**Extended Data Fig. 5d, e**). This data suggests that osimertinib treatment leads to increased expression of *METTL7A* rather than selection of a pre-existing, *METTL7A*-high population in untreated tumor cells.

To interrogate the functional role of METTL7A in acquired osimertinib resistance, we knocked out METTL7A in parental PC9 cells and challenged wildtype and METTL7A knockout (KO) cells with a chronic dose of 1 μM osimertinib over the course of 8 weeks. While parental wildtype cells were able to acquire resistance and resume proliferation in the presence of osimertinib after 5 weeks, METTL7A KO cells were unable to exit the DTP stage and instead largely died by 8 weeks of osimertinib treatment (**Fig. 2j, k**), similar to the phenotype we observed upon depletion of major amplified oncogenes in PC9 and HCC827 cells (**Extended Data Fig. 4c, d**). We rescued this phenotype by overexpressing METTL7A in the KO background and confirmed *METTL7A* expression via western blot and RT-qPCR (**Extended Data Fig. 5c**). We observed a moderate yet significant rescue effect (p < 7 x 10^-6^), suggesting that METTL7A is crucial for cells to acquire resistance to osimertinib (**Fig. 2l, m**). To further confirm that the METTL7A KO phenotype was not due to off-target CRISPR-Cas9 effects, we performed shRNA-mediated knockdown of METTL7A in both PC9 and HCC827 parental cells and validated efficient knockdown via RT-qPCR (**Extended Data Fig. 5f**). Similar to METTL7A KO cells, when treated with osimertinib for approximately 5 weeks, shMETTL7A PC9 and HCC827 cells failed to generate osimertinib-resistant clones compared to parental cells infected with a non-targeting shRNA control lentivirus (**Extended Data Fig. 5g, h**). Moreover, there was no significant growth phenotype in drug-naïve cells with *METTL7A* deficiency (**Fig. 2j-k**, p = 0.21; **Extended Data Fig. 5g, h**).

To investigate if METTL7A is required for the maintenance of osimertinib resistance, we performed *METTL7A* depletion via shRNA in osimertinib-resistant PC9 cells (PC9-OR). We first performed cell viability assays in shMETTL7A and shCTL cells to determine whether METTL7A depletion re-sensitizes cells to osimertinib. We performed this in resistant cells taken off osimertinib for the duration of the experiment (approximately 2 weeks) or in PC9-OR cells in which *METTL7A* was knocked down in the presence of osimertinib. In PC9-OR cells taken off osimertinib, we saw a modest yet significant decrease in resistant colonies in shMETTL7A cells compared to shCTL cells (**Extended Data Fig. 5i, j**). In PC9-OR cells kept on osimertinib for the experiment, we saw an even greater decrease in colonies upon *METTL7A* depletion.

Furthermore, we calculated the “resistance index” for both genotypes by dividing the mean percent area of crystal violet staining in +osi cells by the mean percent area in -osi cells. The resistance index was lower for shMETTL7A cells, suggesting that even after normalizing by the number of cells in the -osi condition, there was still an additional effect of *METTL7A* depletion on drug treatment (**Extended Data Fig. 5k**). Together, these results suggest that METTL7A does indeed have a role in the maintenance of resistance, but because the effect is less significant than *METTL7A* depletion in the DTP stage, we believe that METTL7A’s function is likely more important for the acquisition of resistance.

Finally, we sought to determine whether METTL7A is essential for the acquisition of resistance in other oncogene-amplified cancers commonly treated with targeted therapies. To address this, we utilized BT474 breast cancer cells which harbor *HER2* amplification and acquire resistance to the HER2 inhibitor trastuzumab. We depleted *METTL7A* with shRNAs and treated shMETTL7A and shCTL cells with trastuzumab for approximately 2 months. Like our findings in EGFR-driven LUAD, we found that METTL7A depletion significantly impaired the emergence of trastuzumab-resistant clones without affecting cell viability in drug-naïve cells (**Fig. 2n, o**). This finding suggests that METTL7A might mediate drug resistance in other targeted therapy contexts. Together, these data strongly indicate that METTL7A plays a specific and crucial role during the acquisition of osimertinib resistance.

## METTL7A binds to amplified oncogenes and correlates with CTCF binding

We next aimed to address the mechanism through which METTL7A promotes acquired osimertinib resistance. METTL7A has previously been reported to localize to the endoplasmic reticulum (ER) and inner nuclear membrane but not the nuclear interior or chromatin^30,31^. To elucidate the localization of METTL7A in LUAD cells, we performed immunofluorescence in PC9/PC9-OR and HCC827/HCC827-OR cells that express METTL7A fused to MYC-tag and FLAG epitope tags (METTL7A-MYC-FLAG). We performed this experiment in cells expressing tagged METTL7A due to lack of reliable antibodies that detect endogenous METTL7A via immunofluorescence. Consistent with previous reports, we observed strong accumulation of METTL7A in the cytoplasm of PC9 and HCC827 parental cell lines (**Supplementary Videos 1, 3**). However, we observed both cytoplasmic and nuclear localization of METTL7A in osimertinib-resistant cells (**Fig. 3a, b, Supplementary Videos 2, 4, 5**). We quantified this phenotype and found a significant increase in METTL7A foci in the nucleus of resistant cells compared to parental cells (**Fig. 3c**). In addition, we performed subcellular fractionation followed by western blotting and observed increased METTL7A in the nuclear fraction of osimertinib-resistant cells compared to parental cells, which exhibited similar levels of METTL7A in both the membrane and nuclear fractions (**Extended Data Fig. 6a**).

**Fig. 3.**
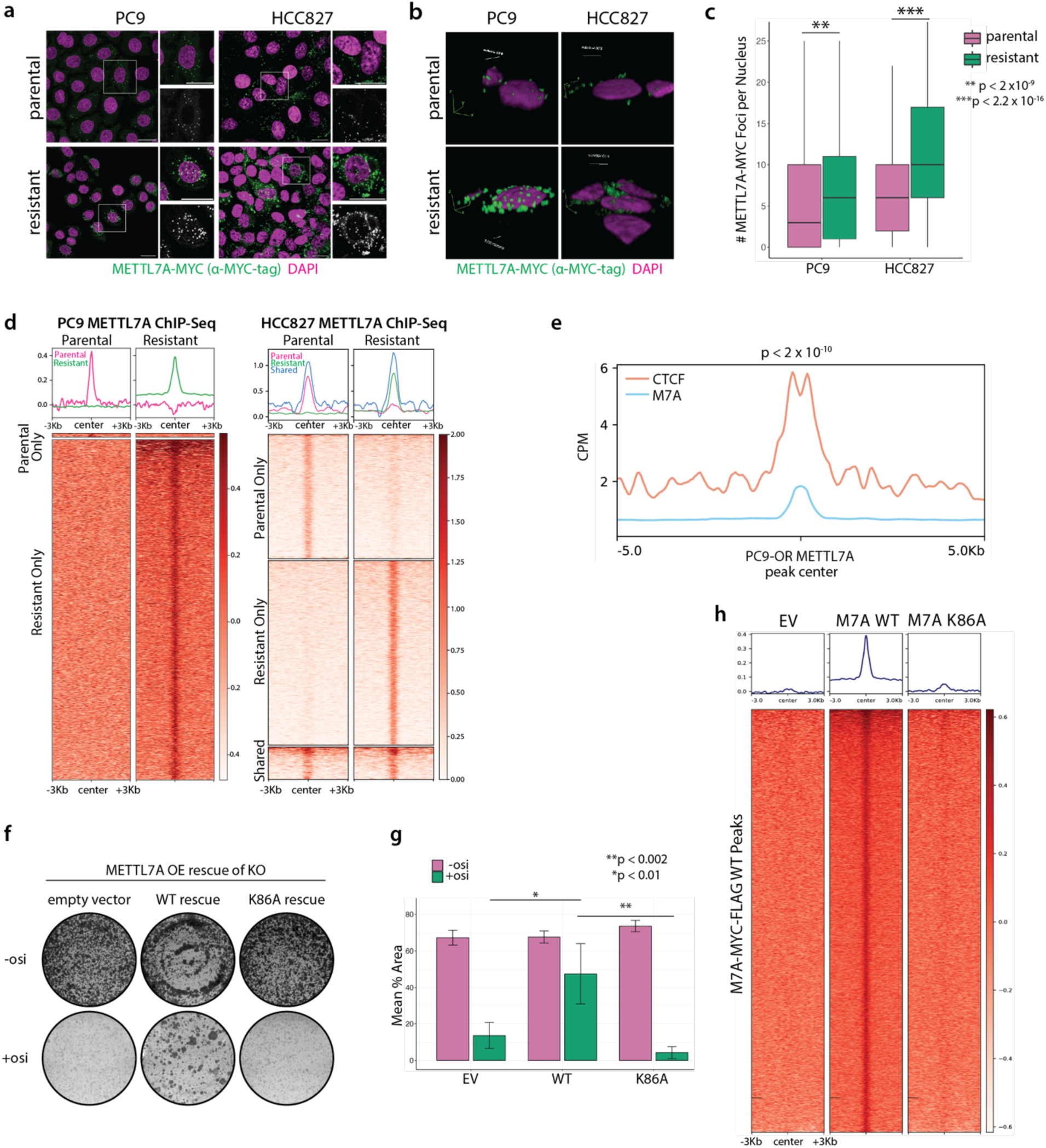
**METTL7A binds to amplified oncogenes**. **a,** Immunofluorescence in PC9 and HCC827 cells expressing MYC-tagged METTL7A shows increased nuclear localization of METTL7A in osimertinib-resistant cells compared to parental cells. Scale bar: 20 µm. **b,** 3D reconstruction of PC9 and HCC827 cells expressing MYC-tagged METTL7A. Scale bar: 5 µm. **c,** Quantification of (a) reveals a significant increase in the number of nuclear METTL7A foci in OR cells compared to parental cells. **d,** Heatmaps of METTL7A MYC-tag ChIP-seq in PC9 and HCC827 cells. Signal is separated based on peaks called in parental cells only, resistant cells only, or peaks present in both parental and resistant cells. Signal is normalized based on CPM and signal from input. **e,** CTCF signal is enriched over METTL7A peaks. Signal is normalized by counts per million (CPM). METTL7A peaks significantly intersect CTCF peaks (p < 2 x 10^-10^, Fisher’s exact test). **f,** Colony formation assays in parental cells treated with or without osimertinib. Colony formation assays were performed in METTL7A KO cells with reconstituted METTL7A WT or K86A (or empty vector control). **g,** Quantification of (f). Error bars represent standard deviation between biological triplicates. Significance determined by unpaired t-test. **h,** MYC-tag ChIP-seq reveals a depletion of signal upon overexpression of M7A-FLAG-MYC K86A mutant, similar to ChIP in cells overexpressing a FLAG-MYC empty vector. Mean ChIP signal between biological replicates is shown. Signal is normalized to input.

Based on the nuclear localization of METTL7A, we wanted to determine whether METTL7A binds chromatin. We performed ChIP-seq in resistant and parental cells overexpressing a METTL7A-MYC-FLAG fusion protein and used a MYC-tag antibody to capture METTL7A’s localization on chromatin. We identified 4,937 and 3,165 high-confidence METTL7A peaks in both PC9-OR and HCC827-OR cells, respectively, that expressed METTL7A-MYC-FLAG fusion protein, compared to very little METTL7A binding in parental cell lines (**Fig. 3d**). To account for the presence of amplicons in OR cells, we were stringent with our peak-calling and called peaks against both ChIP input and IgG samples, thus normalizing for changes in copy number.

Additionally, we normalized our data to PC9-OR and HCC827-OR wildtype cells that did not express the METTL7A-MYC-FLAG fusion protein. Further analysis demonstrated that approximately 60% of amplified oncogenes in both PC9-OR and HCC827-OR cells contained a METTL7A peak (**Extended Data Fig. 6b**). For example, the *RAF1*-containing amplicon locus, which was one of the most amplified loci in PC9-OR cells, contained many METTL7A peaks (**Extended Data Fig. 6c**). We next annotated METTL7A peaks in OR cells and identified an increased proportion of peaks present on promoters compared to the rest of the genome (**Extended Data Fig. 6d**). Additionally, we performed motif and gene ontology analyses on total METTL7A peaks and promoter peaks respectively. While we did not identify any common motifs between PC9-OR and HCC827-OR cells, we did observe CREB1 as a common enriched transcription factor from ChIP Enrichment Analysis (CHEA) and RAD21 from ENCODE (**Extended Data Fig. 6e, f**).

Based on our observation that CTCF is enriched on the boundaries of amplified oncogenes, we next correlated METTL7A binding with PC9 CTCF ChIP. Intriguingly, METTL7A strongly correlated with CTCF signal (**Fig. 3e**), and we observed a significant overlap between METTL7A peaks and CTCF peaks (p < 2 x 10^-10^). As negative controls, we profiled the intersection between METTL7A and H3K9me2 and H3K9me3 deposition based on ChIP-seq performed in parental PC9 cells from the ENCODE Project^25,26^ (ENCSR521GRK and ENCSR555TAX, respectively). Both H3K9me2 and H3K9me3 peaks were significantly excluded from METTL7A peaks, with only 1 H3K9me2 peak intersecting with METTL7A (p < 4 x 10^-6^) and only 27 H3K9me3 peaks intersecting with METTL7A (p < 3 x 10^-7^) (**Extended Data Fig. 6g**). These results further emphasize the specificity of METTL7A binding to CTCF sites. Together, these results suggest that METTL7A binds to amplified genes in osimertinib-resistant cells and correlates with CTCF binding.

We next aimed to determine the critical residues responsible for METTL7A’s chromatin binding activity. We specifically mutated lysine residues which we reasoned might be responsible for METTL7A’s DNA binding activity and honed in on the residue K86. We mutated this residue to alanine (K86A), overexpressed this construct in the *METTL7A* KO background, and validated that mutant METTL7A was expressed at a similar level to WT METTL7A (**Extended Data Fig. 6h**). To test if K86 has a functional role *in vivo*, we treated *METTL7A* KO cells rescued with an empty vector (EV), WT METTL7A, or METTL7A K86A with or without osimertinib and fixed and stained cells after approximately 5 weeks. While METTL7A WT overexpression in *METTL7A*- depleted cells rescued the drug sensitivity phenotype, overexpression of the K86A mutant failed to rescue the phenotype, resulting in significantly fewer resistant clones than KO cells rescued with WT METTL7A (**Fig. 3f, g**). Thus, these results suggest that the K86 residue plays a critical role for METTL7A’s function.

We next tested whether METTL7A K86A affects the localization or chromatin binding activity of METTL7A. We observed a statistically significant decrease in the number of METTL7A nuclear foci in the K86A mutant compared to WT (**Extended Data Fig. 6i, j**). Furthermore, we performed ChIP-seq against MYC-tag in PC9-OR cells overexpressing a FLAG-MYC empty vector (EV); wildtype M7A-MYC-FLAG; or M7A-MYC-FLAG with a K86A mutation. K86A significantly abolished METTL7A’s ability to bind chromatin to a similar extent as cells expressing an empty vector (**Fig. 3h**). Together, these results suggest that the K86 residue is critical for METTL7A’s nuclear localization and binding to chromatin.

To further elucidate the chromatin binding activity of METTL7A, we purified recombinant, MBP-tagged METTL7A from *E. coli* (**Extended Data Fig. 7a**). We incubated MBP-tagged METTL7A with 60 bp dsDNA and ssDNA substrates and ran the protein/DNA complex on a non-denaturing gel. Consistent with our *in vivo* METTL7A binding results, we observed a strong upward gel shift of the MBP-METTL7A-DNA complex at increasing concentrations of METTL7A (**Extended Data Fig. 7b**). We then performed the same assay with the K86A mutant and observed a moderate reduction in binding affinity of the mutant compared to wildtype METTL7A protein (**Extended Data Fig. 7c**). Together, these data strongly support the notion that METTL7A binds to chromatin *in vivo* and to DNA *in vitro*, and the K86 residue may be responsible for METTL7A’s chromatin binding activity.

## METTL7A primes the chromatin landscape for oncogene amplification

Based on the enrichment of CTCF and RAD21 signal over amplicon boundaries, the correlation between METTL7A and CTCF binding, and METTL7A’s binding activity toward DNA, we hypothesized that METTL7A is involved in priming the chromatin landscape during the acquisition of resistance. We performed H3K27ac and RAD21 CUT&RUN in METTL7A wildtype, KO, and rescue cells. We performed this experiment in parental PC9 cells naïve to drug treatment and PC9 cells treated with osimertinib for approximately 6-8 weeks, during which we observed global changes in chromatin architecture but prior to oncogene amplification in the majority of the cells (**Fig. 2b, c, Fig. 4a**). Additionally, this is the time point at which METTL7A is most dynamically upregulated (**Fig. 2i**). Strikingly, our data demonstrated that the global chromatin re-organization was abolished in METTL7A KO cells treated with osimertinib: based on PCA analysis of H3K27ac and RAD21 CUT&RUN signal, the chromatin state in METTL7A KO cells treated with osimertinib was nearly indistinguishable from their drug-naïve counterparts (**Fig. 4b**). Additionally, the peaks gained upon osimertinib treatment in PC9 wildtype cells were those lost upon METTL7A KO in osimertinib-treated cells: 12% of H3K27ac peaks and 55% of RAD21 peaks were lost upon METTL7A KO in osimertinib-treated cells (**Fig. 4c**). We were able to partially rescue this phenotype by overexpressing METTL7A in the KO background (**Extended Data Fig. 8a**). In untreated PC9 cells, we observed a modest decrease in H3K27ac and RAD21 deposition, but this effect was not as strong as in drug-treated cells (**Extended Data Fig. 8b**). This result is consistent with our previous findings that METTL7A is most highly expressed and strongly implicated during the osimertinib resistance acquisition stage rather than in drug-naïve cells. Together, these data suggest that METTL7A is crucial for the chromatin structure changes and epigenetic priming as cells exit the DTP stage but prior to oncogene amplification.

**Fig. 4.**
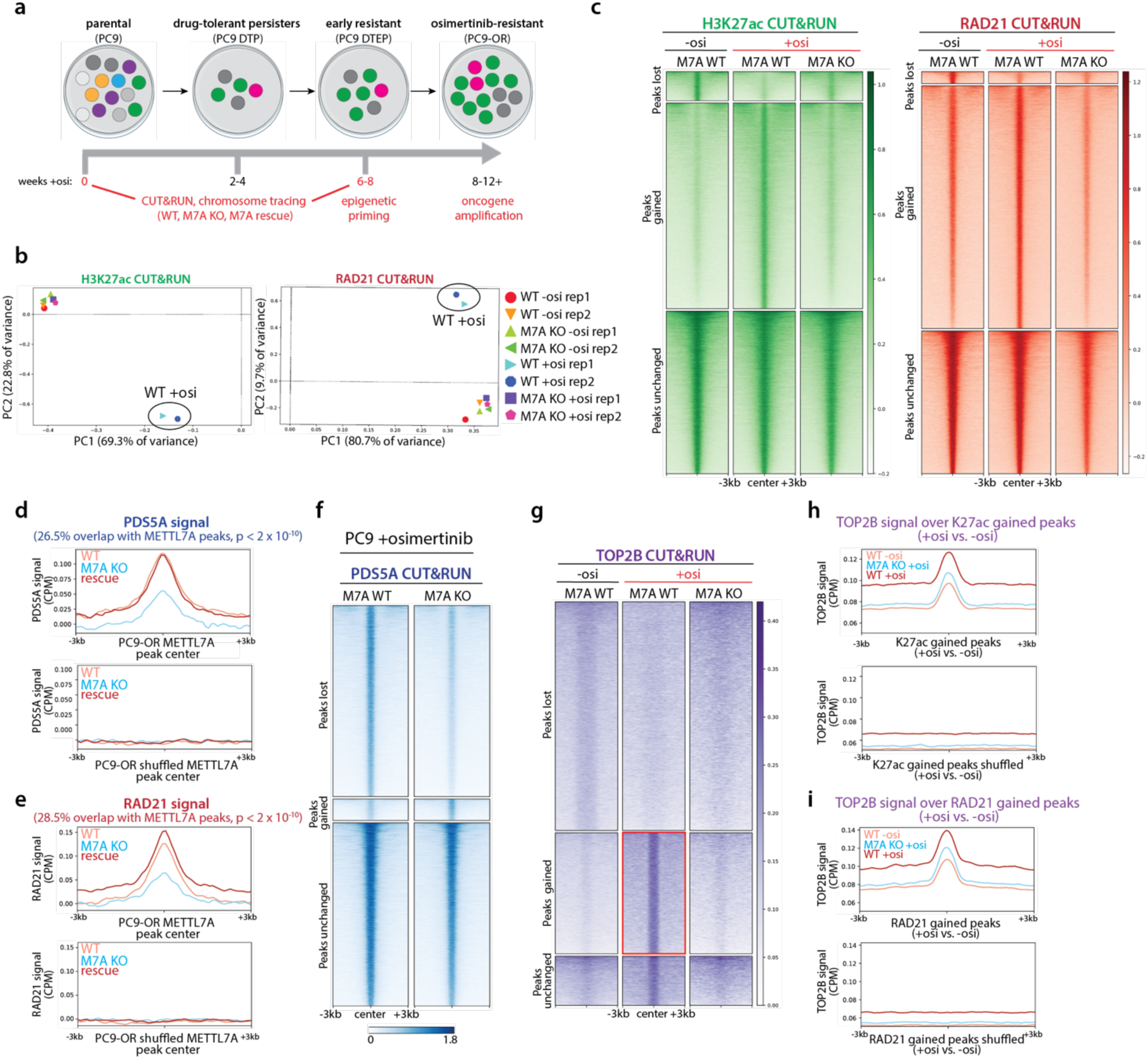
METTL7A primes chromatin via recruitment of cohesin components. **a,** Schematic of PC9 cell line model of acquired osimertinib resistance. Cells emerge from a quiescent, drug-tolerant persister (DTP) state before entering a proliferative resistant state. Time points selected for genomics experiments are highlighted in red. **b,** PCA plots of H3K27ac and RAD21 CUT&RUN bigWig signal shows that METTL7A KO cells treated with osimertinib fail to exit a drug-naïve-like chromatin state. Individual biological replicates are plotted. Signal is normalized based on CPM and H2A CUT&RUN signal is subtracted. **c,** Heatmaps of H3K27ac and RAD21 CUT&RUN signal in PC9 cells treated with and without osimertinib. Signal is centered on peaks that were lost, gained, or unchanged upon osimertinib treatment. Treatment with osimertinib is associated with gain of both H3K27ac and RAD21 peaks. Depletion of METTL7A (M7A) results in reduced H3K27ac and RAD21 binding, resembling the drug-naïve chromatin state. **d,** PDS5A signal is enriched over METTL7A peaks compared to peaks shuffled over the genome. Significance calculated via Fisher’s exact test (p < 2 x 10^-10^). **e,** RAD21 signal is enriched over METTL7A peaks compared to randomly shuffled peaks. Significance calculated via Fisher’s exact test (p < 2 x 10^-10^). **f,** Heatmaps of PDS5A CUT&RUN data in PC9 METTL7A WT and KO cells treated with osimertinib. Signal is centered on peaks that were lost, gained, or unchanged upon METTL7A KO. CUT&RUN signal is normalized by CPM and H2A signal is subtracted. Mean bigWig signal between two biological replicates is shown. **g,** TOP2B CUT&RUN shows an increase in TOP2B intensity in WT cells treated with osi (red box), which is depleted upon *METTL7A* KO. CUT&RUN signal is normalized by *E. coli* spike-in and CPM. Mean bigWig signal between two biological replicates is shown. **h-i,** TOP2B signal is enriched over H3K27ac (h) and RAD21 (i) peaks gained in WT cells treated with osimertinib. TOP2B signal is depleted in *METTL7A* KO cells treated with osi to levels similar to untreated WT cells.

We next wanted to determine the mechanism through which METTL7A primes chromatin. We identified nuclear METTL7A-interacting proteins through nuclear fractionation followed by immunoprecipitation and mass spectrometry in PC9 parental and osimertinib-resistant cells expressing tagged METTL7A. We used the APOSTL SAINT pipeline^32^ to identify significant interactors between two biological replicates of each sample compared to negative control cells that did not express tagged METTL7A (**Supplementary Table 3**). We were particularly intrigued to identify PDS5A as a METTL7A interactor, since PDS5A is known to load and unload cohesin to chromatin and localizes to CTCF loop anchors in mammalian cells^33^. We validated this interaction by performing immunoprecipitation followed by western blotting, and indeed observed interaction between these two proteins (**Extended Data Fig. 8c**). Additionally, we performed proximity ligation assays (PLAs) to further validate this interaction. We performed PLAs between PDS5A and RAD21, which are known to be in close proximity, as a positive control. We identified a significant increase in METTL7A/PDS5A PLA foci in PC9-OR and HCC827-OR cells that express METTL7A-MYC-FLAG compared to wildtype controls (**Extended Data Fig. 8d, e**).

We next determined whether METTL7A-mediated changes in the chromatin landscape were due to METTL7A’s association with PDS5A. We performed PDS5A CUT&RUN in METTL7A wildtype, KO, and rescue cells at the same aforementioned time points. We first correlated PDS5A CUT&RUN signal with METTL7A ChIP-seq peaks and observed a significant overlap (p < 2 x 10^-10^) between PDS5A and METTL7A peaks (**Fig. 4d**). Furthermore, when we centered PDS5A signal from METTL7A wildtype, KO, and rescue cells on the METTL7A peaks from our METTL7A ChIP-seq experiment, we observed that the decreased PDS5A signal occurred at METTL7A binding sites, and PDS5A signal was fully restored in the rescue cells. This further indicates that METTL7A’s interaction with PDS5A affects its chromatin deposition (**Fig. 4d**).

Similarly, we performed the same analysis with RAD21 signal in METTL7A wildtype, KO, and rescue cells and found that RAD21 was depleted in METTL7A KO cells at METTL7A binding sites, and its binding was almost completely restored in the rescue cells (**Fig. 4e**). This further indicates that METTL7A plays a critical role in recruiting the cohesin complex to METTL7A- centered loci. Based on this correlation, we hypothesized that METTL7A affects cohesin deposition via PDS5A recruitment, either directly or indirectly, to chromatin. To address this, we next examined the effects of METTL7A KO on global PDS5A deposition. Indeed, we observed a 47% decrease in PDS5A peaks in METTL7A KO cells compared to wildtype cells (**Fig. 4f**), and PDS5A signal was decreased globally (**Extended Data Fig. 8f**). Similar to our H3K27ac and RAD21 CUT&RUN data, we observed only modest effects of METTL7A KO on PDS5A deposition in parental cells (**Extended Data Fig. 8g**). While PDS5A has been shown to both load and unload the cohesin complex^33–36^, the concomitant loss of both PDS5A and RAD21 binding upon METTL7A KO suggests that PDS5A is likely more responsible for loading of the cohesin complex in this context. In support of this, we observed a strong overlap between both PDS5A and RAD21 peaks in our CUT&RUN data (**Extended Data Fig. 8i**). Furthermore, we ruled out the possibility that differences in PDS5A and RAD21 deposition were due to changes in expression upon METTL7A KO. Indeed, when we probed for total PDS5A and RAD21 protein levels in METTL7A wildtype, KO and rescue cells, we observed no differences in PDS5A and RAD21 expression between the genotypes (**Extended Data Fig. 8j**).

Based on our findings that METTL7A promotes acquisition of osimertinib resistance via PDS5A- mediated cohesin deposition and global chromatin reorganization, we next asked which specific gene targets METTL7A regulates. We observed that in PC9 cells treated with osimertinib for 6-8 weeks, even prior to any oncogene amplification or transcription upregulation, METTL7A targets PDS5A and RAD21 to future amplified oncogenes such as the *RAF1* locus: in METTL7A KO cells, PDS5A and RAD21 are depleted at these “future amplicons”. These sites further correlate with METTL7A deposition based on the METTL7A ChIP-seq data (**Extended Data Fig. 8h**). The “future amplicons” correlate most strongly with CTCF binding, cohesin deposition, and METTL7A peaks.

Previous work has shown that DNA amplification caused by superhelical tension is regulated by topoisomerase 2B, which relieves topological stress at loop anchors^47^. Thus, we wanted to further probe the mechanism of METTL7A-mediated chromatin priming by assessing whether TOP2B functions downstream of this METTL7A-mediated mechanism. To address this, we performed TOP2B CUT&RUN in parental and osimertinib-treated PC9 WT and METTL7A KO cells to determine whether METTL7A mediates gene amplification via TOP2B. Interestingly, we observed decreased TOP2B signal upon METTL7A depletion (**Fig. 4g**). Specifically, we observed increased TOP2B signal in wildtype cells treated with osimertinib, which was depleted upon METTL7A KO (**Fig. 4g**, red box), reminiscent of the RAD21 and H3K27ac deposition change. Hence, we correlated this data with our H3K27ac and RAD21 CUT&RUN data (**Fig. 4g**, red box). Specifically, we profiled TOP2B signal in PC9 WT cells treated with or without osimertinib, or METTL7A KO cells treated with osimertinib. We correlated this signal with the H3K27ac and RAD21 peaks gained in wildtype cells treated with osimertinib and found that TOP2B signal in PC9 WT cells treated with osimertinib correlates with H3K27ac and RAD21 sites gained upon osimertinib treatment (**Fig. 4h, i**). At these sites, TOP2B signal is reduced upon METTL7A depletion. Together, these results suggest that the METTL7A-mediated chromatin priming phenotype may be further regulated by TOP2B.

Finally, we examined if this METTL7A-mediated mechanism directly regulates oncogene amplification. To address this, we performed *RAF1* DNA FISH in *METTL7A* WT and KO cells exiting the DTP stage. In wildtype cells, we observed a moderate yet statistically significant increase in *RAF1* amplification in cells treated with osimertinib after 6-8 weeks (p < 0.0005). However, we observed no significant increase in *RAF1* copy number in *METTL7A* KO cells treated with osimertinib compared to untreated cells (**Extended Data Fig. 9a, b**). These results suggest that METTL7A facilitates gene amplification as cells acquire resistance to osimertinib. We further probed the effect of METTL7A on copy number by depleting *METTL7A* in PC9-OR cells, which we previously showed re-sensitizes PC9-OR cells to osimertinib (**Extended Data Fig. 5i, j**). We similarly performed *RAF1* DNA FISH and observed a significant decrease in the number of *RAF1* foci in shMETTL7A cells compared to shCTL cells (**Extended Data Fig. 9c, d**). Finally, we performed WGS and AmpliconArchitect analysis in PC9-OR WT and *METTL7A* KO cells and observed a global decrease in gene copy number upon *METTL7A* depletion. Together, these results suggest that METTL7A regulates oncogene copy number.

In summary, these results suggest that METTL7A recruits the cohesin complex to CTCF-enriched loci and “primes” chromatin for future amplification.

## METTL7A is a novel regulator of chromatin compaction

To understand how METTL7A regulates chromatin conformation during the acquisition of osimertinib resistance, we performed imaging-based, single-cell chromatin tracing of chromosome 22 in PC9 cells during the acquisition of osimertinib resistance as previously described^37^. This single-cell resolution technique circumvents the high cell number required for standard Hi-C experiments, which is challenging to obtain when cells are treated with osimertinib, especially in the METTL7A KO background. We leveraged a Chr22 library with probes targeting the center of each TAD along chromosome 22 and performed sequential probe hybridization, imaging, and photobleaching to acquire each TAD’s location within the nucleus^37^ (**Fig. 5a**). We performed this experiment in drug-naïve PC9 cells and PC9 cells treated with osimertinib for 6-8 weeks. Because gene amplification had not yet occurred at this time point, the chromatin tracing was not confounded by multiple copies of the same locus.

**Fig. 5.**
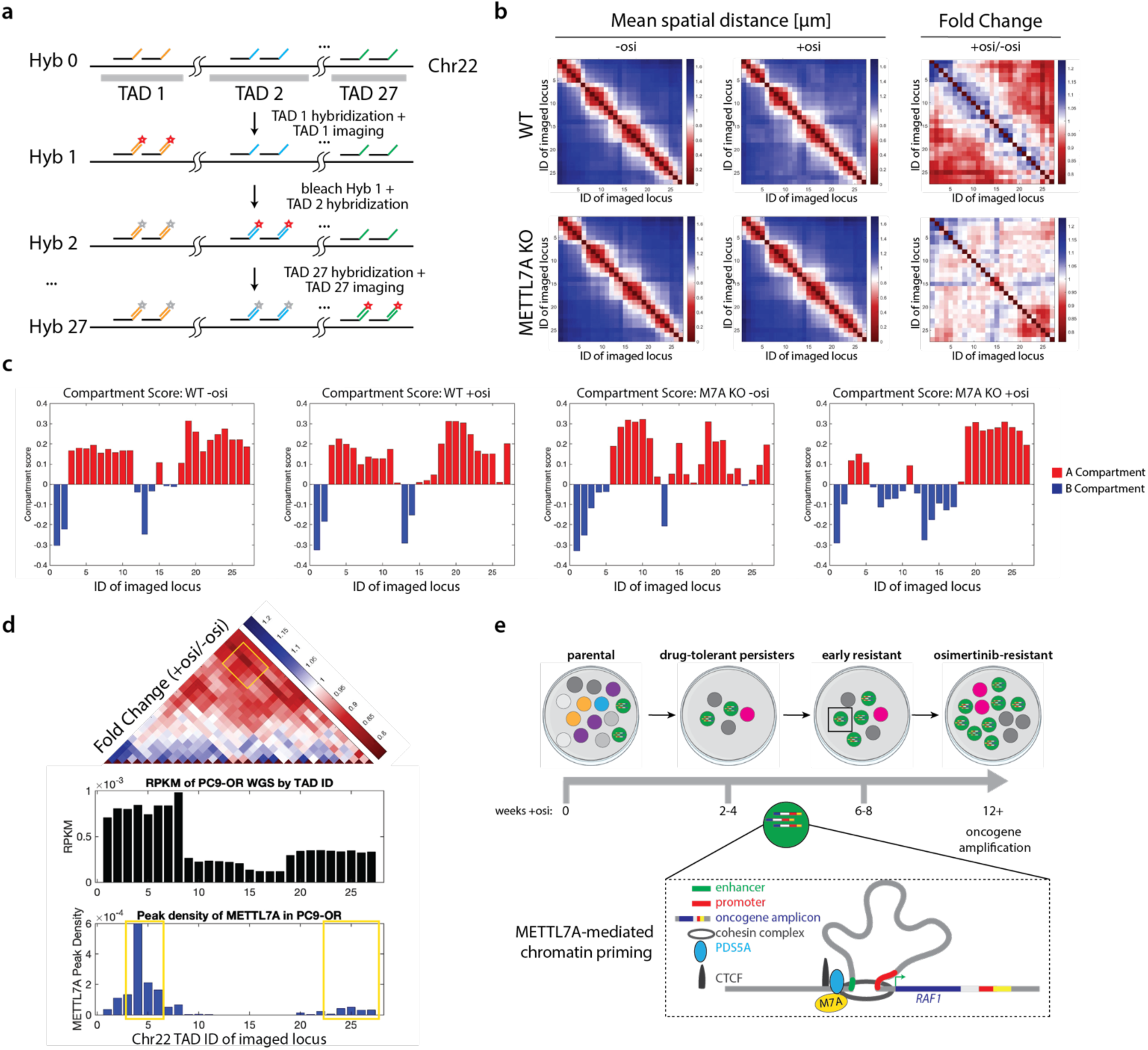
**METTL7A regulates chromatin architecture**. **a,** Schematic of chromosome 22 chromatin tracing. After initial primary probe hybridization to 27 TADs along Chr22, unique secondary probes that bind to primaries for each TAD are sequentially hybridized, imaged, and bleached. **b,** Chr22 tracing reveals increased inter-TAD interaction in wildtype cells treated with osimertinib. In METTL7A KO cells, this gain of inter-TAD contacts is absent. **c,** Compartment score profiles of wildtype and METTL7A KO cells treated with or without osimertinib for 6-8 weeks. **d,** Bar chart showing the WGS coverage (RPKM) in PC9-OR cells within each locus targeted by Chr22 TAD probes and the METTL7A peak density within each TAD based on METTL7A ChIP-seq analysis. **e,** Proposed model of the role of METTL7A during the acquisition of osimertinib resistance. We propose that METTL7A is upregulated as cells acquire resistance to osimertinib. METTL7A localizes to the nucleus, where it binds to amplified oncogenes and affects chromatin architecture via PDS5A-mediated cohesin recruitment, which “primes” these loci for future gene amplification.

By using this method and plotting the mean spatial distance between each TAD along the chromosome, we observed a global decrease in inter-TAD distances in the osimertinib-treated cells compared to parental cells (**Fig. 5b**). This compaction observed in wildtype cells was not present in METTL7A KO cells treated with osimertinib (**Extended Data Fig. 10a**). In contrast, METTL7A KO cells failed to acquire these increased long-range inter-TAD interactions when treated with osimertinib (**Fig. 5b**), supporting our H3K27ac and RAD21 CUT&RUN results revealing that METTL7A KO cells cannot transition into a drug-resistant chromatin state (**Fig. 4b, c**). We further quantified this phenomenon by measuring the number of long-range inter-TAD contact events and found that wildtype cells treated with osimertinib developed a greater frequency of long-range inter-TAD interactions in contrast to METTL7A KO cells (**Extended Data Fig. 10b**). We defined the A and B compartments in these cells and observed that in wild-type cells, osimertinib treatment led to small changes in A and B compartment designation (**Fig. 5c**). However, loss of METTL7A led to significant changes in compartment scores in both untreated and osimertinib-treated KO cells compared to wildtype cells (**Fig. 5c**). Using these A and B compartment definitions, we quantified the long-range AA, AB, and BB inter-TAD contact frequencies and specifically observed that compared to untreated cells, osimertinib-treated cells had a significant increase in long-range AA and AB compartment interactions whereas METTL7A KO osimertinib-treated cells failed to develop such long-range interactions (**Extended Data Fig. 10c**).

Based on our findings that METTL7A binds to amplified loci in resistant cells and affects chromatin architecture even before oncogene amplification has occurred, we next wanted to address whether there was a correlation between the decrease of inter-TAD distance in wildtype cells treated with osimertinib, METTL7A binding based on our previous ChIP-seq analysis, and the amplification of genomic loci in resistant cells. To address this, we plotted Chr22 tracing data alongside our METTL7A ChIP-seq data and whole-genome sequencing. We first observed that the highest fold change between drug-treated cells and parental cells occurred between TADs 1-10 and TADs 23-25. These regions are of particular interest because these are the loci that eventually become amplified in PC9-OR cells (**Fig. 5d**). Additionally, the increase of inter-TAD proximity in wildtype cells treated with osimertinib occurred at loci containing the highest density of METTL7A peaks based on our ChIP-seq analysis (**Fig. 5d**). For example, TADs 3-6 and 24- 27 contained the highest density of METTL7A peaks, and these TADs were also the regions that displayed the greatest increase in spatial proximity in wildtype cells treated with osimertinib (**Fig. 5d**). Together, these data suggest that METTL7A promotes long-range inter-TAD interactions during the acquisition of osimertinib resistance, which may be permissive to the formation and evolution of complex oncogene amplicons.

Based on these data, we propose that METTL7A promotes osimertinib resistance in EGFR-mutant LUAD by binding to and remodeling chromatin architecture. Specifically, as cells are acquiring resistance to osimertinib, METTL7A localizes to the nucleus, where it binds to oncogenes. Through the recruitment of PDS5A and the cohesin complex, METTL7A has a chromatin priming effect, where it remodels chromatin architecture to facilitate the future amplification of these oncogenes (**Fig. 5e**). Because METTL7A plays a critical role during the acquisition of osimertinib resistance – specifically at the DTP exit stage – these findings have the potential to lead to novel therapeutic strategies to prevent osimertinib resistance from occurring in patients.

## Discussion

Acquired resistance to TKIs is a major clinical challenge in the treatment of EGFR-mutant LUAD. While it has long been established that oncogene amplification is a common feature of osimertinib-resistant LUAD tumors, the mechanisms that promote gene amplification are less understood. Here, we identified a unique chromatin signature of drug-naïve tumors that correlates with the boundaries of oncogene amplicons in drug-resistant tumors, which exhibit diverse amplicon structures. Specifically, we found that CTCF is enriched in drug-naïve tumors over the boundaries of “future amplicons”, with a biased enrichment over one side of the future amplicon. This finding builds on recent work showing that CTCF deposition affects the copy number and rearrangement of the *KMT2A* gene amplification in leukemia^24^. While the importance of 3D chromatin organization in driving oncogene expression in cancer has previously been observed^19^, our findings could have broader implications for the identification of loci that are especially susceptible to oncogene amplification.

Furthermore, we have identified a novel function of the protein METTL7A in regulating this chromatin landscape. Specifically, we have found that METTL7A localizes to the nucleus of osimertinib-resistant LUAD cells, where it binds to chromatin to facilitate cohesin-mediated chromatin reorganization via interacting with the cohesin-associated factor PDS5A. When METTL7A is depleted from parental LUAD cells, cells cannot transition into a drug-resistant chromatin state, which we propose is due in part to decreased long-range inter-TAD chromatin contacts required to form highly complex oncogene amplicons demarcated by CTCF and cohesin. Together, this phenotype in METTL7A-depleted cells prevents the acquisition of osimertinib-resistant LUAD clones.

During the acquisition of TKI resistance, cells transition through a quiescent, drug-tolerant persister (DTP) stage, marked by a defined, reversible, epigenetic state. We found that METTL7A was specifically upregulated during this time point, before the onset of oncogene amplification. In parental cells lacking METTL7A, we did not notice any cell viability defects, and instead, only observed impaired cell viability upon treatment with osimertinib. Furthermore, we identified that METTL7A promotes trastuzumab resistance in *HER2*-amplified BT474 breast cancer cells, which may suggest that METTL7A may more broadly promotes resistance to targeted therapies in other oncogene-amplified cancers. Future work should expand this finding to additional oncogene-amplified models and determine whether *METTL7A* depletion has a phenotype in cancers that do not utilize oncogene amplification as a mechanism of resistance.

While the current work primarily focused on the acquisition of resistance rather than tumors that are already resistant to osimertinib, it is important to consider METTL7A’s potential function in maintaining the drug-resistant state. We observed a moderate yet significant effect of *METTL7A* depletion on the re-sensitization of resistant cells to osimertinib, suggesting that METTL7A is a dependency of osimertinib-resistant cells. Furthermore, we observed a decrease in oncogene copy number in *METTL7A*-depleted, resistant cells, which further hints at METTL7A’s potential function in maintaining resistance. However, it is possible that METTL7A’s function in maintaining drug resistance may be decoupled from its role in regulating oncogene amplification, since previous work has shown that *RAF1* knockdown or MEK inhibition in PC9- OR cells harboring *RAF1* amplification does not re-sensitize resistant cells to osimertinib^15^.

Thus, future work should examine how and whether METTL7A regulates drug resistance in tumors that have already acquired resistance.

METTL7A has previously been observed to localize to the endoplasmic reticulum and the inner nuclear membrane, and thus a nuclear or chromatin function of this protein has not previously been uncovered. Previous work has suggested that METTL7A has methyltransferase activities toward lncRNA^38^ and thiol group^39^ substrates. While we cannot rule out the possibility that these additional functions of METTL7A may contribute to osimertinib resistance, our work nevertheless has identified a completely novel, chromatin and DNA-binding function of this relatively understudied protein. Future biochemical and structural work will investigate whether METTL7A has additional methyltransferase activity toward chromatin and whether its chromatin binding activity is distinct from its methyltransferase function. Here, we identify an additional residue, K86, that is crucial for METTL7A’s function and DNA binding activity. Specifically, we have found that mutating this key residue impairs the ability of METTL7A to rescue the knockout phenotype. Furthermore, this mutant abolishes METTL7A’s DNA binding activity. While we did not explore the putative methyltransferase function of METTL7A in this context, it is important to point out that other groups have shown METTL7A has catalytic activity toward substrates such as lncRNAs^38^ and thiol-containing small molecules^39,48^. Additionally, METTL7A has homology to METTL7B, which has also been shown to display activity toward small molecules and RNA^48,49^. Intriguingly, we observed that METTL7B expression was significantly decreased in osimertinib-resistant tumors. This inverse relationship between METTL7A and METTL7B could suggest that the two proteins have redundant functions, and thus, tumors only need to upregulate one to acquire resistance. While we did not perform any mechanistic studies with METTL7B, it would be interesting to further explore its potential role in other models of drug resistance.

In summary, we have identified a unique epigenetic priming mechanism regulated by METTL7A that occurs during the acquisition of TKI resistance in EGFR-mutant LUAD to drive oncogene amplification. We believe that this work is novel at both the mechanistic and translational levels: METTL7A not only displays a unique chromatin binding and priming function, but it also lays the groundwork for the development of therapies that could specifically prevent TKI resistance. Future work will further investigate the possibility of targeting METTL7A in LUAD and other tumor types.

## Methods Cell culture

PC9 cells were obtained from ATCC, H1975 and HCC827 cells were a gift from K. Politi, and BT474 cells were a gift from Qin Yan. PC9, H1975, HCC827 and BT474 cells were cultured in RPMI-1640 medium (Thermo Fisher, 11875119) with 1% penicillin-streptomycin (pen-strep, Thermo Fisher, 15140122) and 10% fetal bovine serum (FBS). HEK 293T cells were obtained from ATCC (CRL-3216) and maintained in DMEM-F12 (Thermo Fisher, 11330057) supplemented with 1% pen-strep and 10% FBS. All cells were passaged with 0.25% trypsin (Thermo Fisher, 25200056) after reaching 70-90% confluency. PC9-OR, H1975-OR and HCC827-OR were generated by treating parental sensitive cells with escalating doses of osimertinib (Selleckchem, S7297) over the course of 12 weeks. Resistant cells were subsequently maintained in 1 uM osimertinib. All cell lines were maintained at 37°C with 5% CO2. All cell lines were tested regularly for mycoplasma at the Yale Virology Core.

## Colony formation assay

For the colony formation assays performed in **Fig. 2j**, PC9 cells were seeded onto 6-well plates in triplicate with 1000 cells seeded per well. For all other colony formation assays, 20,000 cells were seeded onto each well of a 6-well plate in triplicate. In both cases, one day after seeding, PC9 and HCC827 cells were treated with either 1 uM or 0.1 uM osimertinib, respectively.

HCC827 cells were treated with a lower concentration of osimertinib due to their increased sensitivity to drug compared to PC9 cells. Media was changed every 3 days. Osimertinib-treated cells were fixed at the indicated time points with 100% cold methanol for 10 min at -20°C. Untreated control cells were fixed once cells were confluent. Cells were subsequently stained with crystal violet (0.5% crystal violet in 25% methanol) for 10 min at room temperature and washed with diH2O. Plates were imaged at 800 dpi as TIF files. Green channel of each image was isolated and the area of each well was quantified using ImageJ. Three biological replicates were performed for each experiment.

For the colony formation assay performed with BT474 cells, 3,000 cells were seeded per well in technical and biological triplicates. One day after seeding, trastuzumab was added at a concentration of 10 μg/mL. Media was replaced every 2 days for 2 months. Cells were fixed, stained, and imaged as described for the LUAD cells.

## Cell viability assay

Cells were seeded onto 96-well plates in technical triplicate with 1000 cells (PC9 and H1975) or 1500 cells (HCC827) per well. 24 hours after seeding, osimertinib was added at the indicated concentrations. Four days after seeding, cells were fixed for 15 min in 4% formaldehyde and washed two times with PBS. Hoechst (Thermo Fisher, H3570) was added at a final concentration of 1 ug/mL. Plates were imaged using Cytation 3 (BioTek), and CellProfiler was used to quantify the number of cells per well based on Hoechst staining. Percent viability was calculated by normalizing to the lowest concentration of osimertinib (100% viability). Three biological replicates were performed for each experiment.

## Lentivirus production

293T cells were seeded into 10 cm dishes. Once cells reached 80-90% confluency, cells were transfected with 10 ug packaging plasmid (psPAX2, Addgene 12260), 2 ug envelope plasmid (PMD2.6, Addgene 12259) and 10 ug of expression vector using the jetPRIME™ DNA and siRNA Transfection Reagent (Polyplus) according to the manufacturer’s instructions. Virus was collected 48-72 hours post-transfection and filtered through a 0.45 um filter. Virus was concentrated using the Lenti-X Concentrator kit (Takara Biosciences) according to the manufacturer’s instructions and resuspended in PBS.

## Generation of *METTL7A* KO clones

*METTL7A* guide RNAs were designed using the Benchling CRISPR tool. sgRNAs were designed to target the following two loci (5’ è 3’): TGTGATATACAACGAACAGA and CCTCGCGGAGAATCCGCTCC. Single gRNAs were cloned into a lentiviral CRISPR Cas9 plasmid (Addgene 52963) using the BbsI restriction digest method from the Zheng Lab^40^. Lentivirus was prepared as previously described. Cells were transduced with lentivirus and selected for 10-14 days with puromycin (Thermo Fisher A1113803). Cells were then single-cell sorted into 96-well plates using a FACS AriaII. Individual clones were screened for homozygous knock out via genotyping using primers METTL7A_GT_F and METTL7A_GT_R, Sanger sequencing, and RT-qPCR for *METTL7A* using primers METTL7A_RNA_F, METTL7A_RNA_R, actin-F, and actin-R (**Supplementary Table 4**).

## shRNA-mediated gene knockdown

shRNA-mediated gene knockdown cells were created by lentiviral transduction using the PLKO.1 backbone. shRNA lentiviruses were produced by the Yale Functional Genomics Core facility by co-transfecting HEK293T cells with packaging vectors psPAX2 (Addgene plasmid #12260) and pMD2.G (Addgene plasmid #12259) together with lentiviral transfer constructs. Viral supernatant was collected 48h and 72h after transfection and filtered with a 0.45-µm filter. Cells were infected with filtered viral supernatant diluted 1:5 in growth media. Two days after infection, cells were selected with 1 ug/mL puromycin (Thermo Fisher A1113803). Selection was performed for at least 10 days before downstream analysis. The following shRNAs from the The RNAi Consortium (TRC) were used: TRCN0000197115 (RAF1), TRCN0000195502 (RAF1), TRCN0000121248 (MET), TRCN0000121233 (MET), TRCN0000240738 (METTL7A) and SHC002 (non-targeting control).

## DNA FISH probe design

Oligopaint FISH probe libraries were constructed as previously described^41^. In brief, each oligo is a size of 100-nucleotides consisting of a 30-nucleotide (nt) homology to the hg19 genome. Each library subpool consists of a unique set of primers for PCR amplification. The forward primer consists of a 30-nt T7 promoter sequence for in vitro transcription followed by a 20-nt forward primer sequence. A unique secondary 30-nt sequence for each library subpool was appended before the reverse primer sequence. Individual FISH probes were generated by PCR amplification, in vitro transcription and reverse transcription. DNA secondary probes that were conjugated with either Alexa-488, Alexa-594, or Alexa-647 were used in FISH experiments to the unique secondary sequence for each library subpool.

The Oligopaint-covered genomic regions (hg19) used in this study are as follows: chr3: 12,621,717-12,721,700 (hg19_RAF1_100kb), chr3: hg19: 20,505,612-22,005,610 (hg19_Landmark_Chr3_1.5Mb), chr20: 58,425,276-58,525,044 (hg19_SYCP2_100kb), chr20: chr20: 51,500,001-53,000,000 (hg19_Landmark_Chr20_1.5Mb), chr8: 128,730,038-128,833,605 (hg19_MYC_100kb), chr8: 116,860,000-118,680,000 (hg19_Landmark_Chr8_1.5Mb), chr7: 116,350,000-116,450,000 (hg19_MET_100kb), chr7: 55,135,000-55,235,000 (hg19_EGFR_100kb), chr7: 89,500,000-91,000,000 (hg19_Landmark_Chr7_1.5Mb), chr3: hg19: 12,366,001-12,866,000 (hg19_RAF1_500kb). ssDNA oligo pools were ordered and synthesized from GenScript. Secondary FISH probes conjugated with fluorophores were ordered and synthesized from Integrated DNA Technologies.

## Interphase DNA FISH

Cells were fixed in 4% PFA diluted in 1x PBS for 10 minutes. Cells were then washed with PBS and permeabilized in 0.5% Triton-X100 for 30 min. Cells were then washed with 1x PBS twice followed with 0.1M HCl treatment for five minutes. Samples were then treated with 20ul of RNase A/T1 (ThermoFisher, EN0551) in 1mL PBS for 40 minutes at 37C. Cells were washed two times with 2x SSC followed by a 30-minute incubation at room temperature in 2x SSC + 0.1% Tween20 (2x SSCT) + 50% (v/v) formamide (Millipore Sigma, F9037). Fresh hybridization buffer was created consisting of 2x SSC, 50% formamide, and 20% dextran sulfate (EMD Millipore, S4030). 200ng of FISH probes were added to 10ul of hybridization buffer, and the coverslip was denatured at 86°C for 5 min. Samples were incubated with primary probe at 37 °C for 16 h in a humidified chamber. Samples were then washed twice for 15 min in pre-warmed 2x SSCT at 60 °C and then were further incubated at 2x SSCT for 15 min at room temperature.

Samples were then incubated with secondary probes (2 uM fluorescently-conjugated secondary probe diluted in 30% formamide + 2x SSC) for 30 min at room temperature. Coverslips were washed with 40% formamide in 2x SSC for 5 minutes followed by additional washes in 2x SSC. Coverslips were then mounted in Prolong Gold Antifade Mountant with DAPI. CellProfiler was used to quantify interphase FISH images.

## Metaphase DNA FISH

Cells in metaphase were prepared by KaryoMAX (Gibco) treatment at 0.1 μg/mL for 3 h. Single-cell suspension was then collected and washed by PBS, and treated with 75 mM KCl for 15 min. Samples were then fixed by 3:1 methanol:glacial acetic acid, v/v and washed once with the fixative. Samples were stored at -20C before use. A slide was heated to 86C for one minute before dropping the cell suspension onto the slide. Cells were then dropped onto the slide and heated for one minute at 86°C. Slides were equilibrated with 2x SSC for five minutes and DNA FISH was performed using the same protocol as interphase DNA FISH.

## Microscopy

Images were acquired on a Leica SP5, Zeiss LSM 880, or Leica Stellaris at either 40x or 63x for interphase FISH samples. For metaphase FISH samples, images were acquired at 100x. All images were taken as z-stacks at either 0.5 um steps or optimized steps depending on the pinhole size. Z-stack images were further processed as maximum intensity projection images in ImageJ/FIJI and analyzed in CellProfiler or ImageJ/FIJI.

## Whole genome sequencing

Library preparation and sequencing was performed by Novogene. Briefly, 1 ug of genomic DNA was randomly sheared, and fragments were end-repaired, A-tailed, and ligated with Illumina adapters. Size selection was performed, followed by PCR amplification and purification. The resulting library was quantified with Qubit and RT-PCR, and bioanalyzer was used to determine the size distribution of the library. Libraries were then pooled and loaded onto an S4 flowcell and sequenced with NovaSeq 6000 with 10x genomic coverage per sample.

## AmpliconArchitect and AmpliconClassifier

WGS FASTQ files were processed using the AmpliconSuite-pipeline v0.1555.2 to call AmpliconArchitect v1.3.r5 and AmpliconClassifier v0.5.3. and CNVKit v0.9.10against GRCh38. Default settings were run with GRCh38 set as reference genome. Amplicon complexity scores were retrieved from the output of AmpliconClassifier and measures the diversity of amplicons. To perform grouped analysis in paired sensitive and resistant cells, GroupedAnalysis.py was run to collate all seed regions before invoking AmpliconArchitect for paired parental and osimertinib-resistant cell lines.

## PDX Patient Sample Processing

Viable tumors or single-cell suspensions of PDXs were provided by the NCI CAPR, which were derived from tumors that progressed on osimertinib treatment from NCT02759835. DNA and RNA was isolated from approximately 30mg of tissue using the All Prep DNA/RNA Mini Kit (QIAGEN).

## WES Analysis of Oncogene Amplification

BAM files were retrieved from dbGaP accession code phs002001. CNVKit v0.9.9 was used to analyze for copy number variation between post-osimertinib treated tumors and pre-osimertinib tumors. The pre-treated sample was inputted using the --normal flag. The --target flag was provided from Agilent as S06588914_Regions.bed file as the original patient samples were processed using SureSelect Clinical Research Exome Kit. The --annotate flag was provided from UCSC as refFlat.txt for hg19. RPKM for RNA-seq were retrieved from the supplemental information of Roper et al. To produce the graphs in Extended Data Fig. 2, amplified oncogenes with a cutoff provided in **Supplementary Table 2** were used. Oncogenes were retrieved from the supplemental information of Luebeck et al^16^.

## RNA sequencing

RNA was extracted using the RNeasy Mini Kit (Qiagen, 74104O) and concentrations were measured via Nanodrop. Library preparation was performed using approximately 1 ug of RNA. Library preparation and sequencing were performed by Novogene. Briefly, mRNA was purified from total RNA with poly-T oligo-attached magnetic beads. Fragmentation was performed and the first strand cDNA was synthesized with random hexamer primers followed by second strand cDNA synthesis. cDNA was then subject to end repair, A-tailing, adapter ligation, size selection, PCR amplification and purification. Qubit and RT-PCR were used to quantify resulting libraries, and bioanalyzer analysis was performed to determine size distribution. Libraries were pooled and loaded onto an S4 flowcell and sequenced with NovoSeq 6000.

## RNA-seq analysis

Sequencing reads were aligned with STAR version 2.7.7a using a GRCh38 STAR index. Gene expression was quantified using featureCounts from Subread version 2.0.3. DESeq2 was used for differential analysis. To profile differentially expressed putative and known epigenetic factors, the GSEA database was used to search for epigenetic gene lists. A single list of known or putative epigenetic and/or chromatin factors was curated by combining GSEA lists: “REACTOME_EPIGENETIC_REGULATION_OF_GENE_EXPRESSION,” “GOMF_TRANSFERASE_ACTIVITY,” “GOMF_DOUBLE_STRANDED_METHYLATED_DNA_BINDING”, and “GOBP_NEGATIVE_REGULATION_OF_GENE_EXPRESSION_EPIGENETIC”.

## ChIP-Sequencing

ChIP-seq was performed using the Active Motif ChIP-IT Express Kit (Active Motif 53008) according to the manufacturer’s instructions. Approximately 2 x 10^7^ cells were used per biological replicate. Chromatin was sheared using a Covaris S220 using a peak power of 175W, duty factor of 10%, 200 cycles/burst, and 160 seconds per run. One uL of de-crosslinked chromatin was analyzed via Agilent TapeStation to determine fragment size, and DNA concentration was measured using Qubit (Invitrogen Q32854). The following antibodies were used: IgG isotype control (CST 5415S) and c-MYC-tag (Cell Signaling Technologies 2276S). 20 ug of chromatin was pre-incubated with antibody overnight at 4°C. Protein G beads were subsequently added to antibody-bound chromatin and incubated for 4 hours at 4°C. 2.5 ng of immunoprecipitated chromatin was subject to library preparation using EpiCypher CUT&RUN Library Kit (EpiCypher 141001) according to the manufacturer’s instructions. Pooled libraries were sequenced by Novogene with NovaSeq 6000 with 20 million paired-end reads sequenced per library.

## ChIP-seq analysis

Adapter sequences from paired-end sequencing reads were trimmed with Trimmomatic and quality control was performed using FastQC. Bowtie2 version 2.4.2 was used to align reads to the GRCh37 genome using the –very-sensitive flag and allowing for one mismatch. SAMtools version 1.16 was used to remove unmapped reads, secondary alignment reads, and reads with a Phred score of <30. Macs2 version 2.2.7.1 was used to call peaks with default peak-calling settings. Peaks were called against input chromatin. Peaks from 2 biological replicates were merged. Deeptools version 3.5.1 was used for ChIP-seq data visualization. Plots depicting ChIP-seq signal were generated by using Deeptools bamCoverage to normalize signal by CPM and duplicates were ignored. Peak intersections were performed using BEDTools v2.30 and quantified using BEDTools Fisher. The mean bigWig signal between biological replicates was plotted.

## GO Analysis

ChIP-seq peaks were annotated to the nearest gene using ChIPseeker and promoter peaks were filtered before running EnrichR (https://maayanlab.cloud/Enrichr/). The top five significant terms ranked by p-value from ENCODE and ChEA Consensus TFs from ChIP-X were plotted.

## MEME Analysis

FASTA sequences were generated using the center of ChIP-seq peaks with flanking windows of 250bp (total window size of 500bp). Enriched motifs were identified using MEME v5.4.1 meme- chip. The top three STREME outputs ranked by p-value were visualized in R using universtalmotif v3.15.

## CUT&RUN

CUT&RUN was performed using the CUTANA ChIC/CUT&RUN Kit (EpiCypher 141048) according to the manufacturer’s instructions. Approximately 5 x 10^5^ cells were used per reaction. The following antibodies were used: Histone H2A (12349), PDS5A (Thermo Fisher PA5-57755), RAD21 (Active Motif 91245), H3K27ac (CST 8173), TOP2B (abcam ab72334), and IgG (EpiCypher 13-0042). One microliter of chromatin was analyzed via Agilent TapeStation to determine fragment size. DNA concentration was measured using Qubit (Invitrogen Q32854). Library preparation was performed using the CUTANA CUT&RUN Library Prep Kit (EpiCypher 14-1001) using 2.5 ng chromatin as input. Pooled libraries were sequenced by Novogene with NovaSeq with 20 million paired-end reads sequenced per library.

## CUT&RUN analysis

Adapter sequences from paired-end sequencing reads were trimmed with Trimmomatic and quality control was performed using FastQC. Bowtie2 version 2.4.2 was used to align reads to the GRCh37 genome using the –very-sensitive flag and allowing for one mismatch. SAMtools version 1.16 was used to remove unmapped reads, secondary alignment reads, and reads with a Phred score of <30. Macs2 version 2.2.7.1 was used to call peaks with default peak-calling settings. Peaks were called against H2A. IgG peaks were subtracted from BED files using BEDTools v2.30. IDR was used to determine significant peaks between two biological replicates. Deeptools version 3.5.1 was used for data visualization. Plots depicting CUT&RUN signal were generated by using Deeptools bamCompare to normalize signal by CPM and to H2A, and duplicates were ignored. In all plots, the mean normalized bigWig signal between two biological replicates is plotted. Peak intersections were performed using BEDTools v2.30 and quantified with BEDTools Fisher.

For TOP2B CUT&RUN, due to higher levels of noise in the sequencing data, SEACR was used to call peaks. bigWig signal was normalized via *E. coli* spike-in DNA, and normalized bigWig signals were used as inputs for SEACR using the “stringent” settings. BEDTools was used to intersect peaks between biological replicates. Downstream visualization with Deeptools was performed as previously described.

## Subcellular fractionation and protein immunoprecipitation

Approximately 2 x 10^7^ wildtype and METTL7A-MYC-FLAG-tag-expressing cells were subject to subcellular fractionation using the Subcellular Fractionation Kit for Cultured Cells (Thermo Fisher 78840) according to the manufacturer’s instructions. The protein concentration of each fraction was measured with a BCA assay (Pierce 23225). Anti-FLAG magnetic beads (Millipore M8823) were washed 3x with TBS buffer (50 mM Tris-HCl pH 7.4, 150 mM NaCl), and 30 uL of bead slurry was added to 500 ug of nuclear lysate diluted in 1 mL TBS. Nuclear lysates were incubated with beads for 2 hours at 4°C. Beads were spun down and subsequently washed 3x 10 min with TBS buffer at 4°C. Immunoprecipitated proteins were eluted by incubating beads 2x 30 min with 150 ng/uL FLAG peptide at 4°C. Eluted proteins were sent for mass spectrometry analysis. Two biological replicates were performed for each sample.

## Mass spectrometry

Mass spectrometry was performed by the IDeA National Resource for Quantitative Proteomics in Little Rock, AR, with the following procedure: Protein samples were reduced, alkylated, and digested using filter-aided sample preparation^42^ with sequencing grade modified porcine trypsin (Promega). Tryptic peptides were then separated by reverse phase XSelect CSH C18 2.5 um resin (Waters) on an in-line 150 x 0.075 mm column using an UltiMate 3000 RSLCnano system (Thermo). Peptides were eluted using a 60 min gradient from 98:2 to 65:35 buffer A:B ratio (Buffer A = 0.1% formic acid, 0.5% acetonitrile; Buffer B = 0.1% formic acid, 99.9% acetonitrile). Eluted peptides were ionized by electrospray (2.4 kV) followed by mass spectrometric analysis on an Orbitrap Eclipse Tribrid mass spectrometer (Thermo). MS data were acquired using the FTMS analyzer in profile mode at a resolution of 120,000 over a range of 375 to 1200 m/z.

Following HCD activation, MS/MS data were acquired using the ion trap analyzer in centroid mode and normal mass range with a normalized collision energy of 30%. Proteins were identified by database search using MaxQuant (Max Planck Institute) with a parent ion tolerance of 3 ppm and a fragment ion tolerance of 0.5 Da. Scaffold Q+S (Proteome Software) was used to verify MS/MS based peptide and protein identifications. Protein identifications were accepted if they could be established with less than 1.0% false discovery and contained at least 2 identified peptides. Protein probabilities were assigned by the Protein Prophet algorithm^43^.

## Mass spectrometry analysis

Analysis of mass spectrometry hits was performed with the SAINTExpress/APOSTL pipeline. A samples report was generated from Scaffold using the following parameters: protein threshold of 99%, minimum number of peptides = 1, and peptide threshold of 95%. Known contaminants such as keratin were manually removed. The samples report containing both biological replicates per sample was then uploaded to the APOSTL Galaxy Server. Wildtype cells not expressing the METTL7A-MYC-FLAG fusion protein were set as negative controls. The CRAPome was queried to exclude known contaminants from analysis. Thresholds were set by using a SAINT score > 0.5 and log_2_FC > 2.

## IP/western blot validation of mass spectrometry

To validate significant interactors from mass spectrometry, nuclear lysates were prepared via the Dignam and Roeder method^44^. Nuclear lysates were quantified with BCA assay, and 300 ug protein was diluted in 400 uL PBS with protease inhibitors. Lysates were pre-incubated with antibodies overnight at 4°C. The following antibodies were used: IgG (CST 5415S), FLAG (Sigma F1804), and MYC-tag (CST 2276S). Dynabeads™ M-280 Sheep Anti-Mouse IgG (Invitrogen 11201D) were washed 3x with PBS + 0.1% BSA, and 50 uL bead slurry were added to each antibody/lysate mixture. Lysates were incubated with beads for 2 hours at 4°C. Beads were washed 3x 5 min with PBS + 0.1% BSA and eluted with 2X Laemmli buffer + BME via boiling for 5 min at 95°C.

## Chromatin tracing

The design of a chromatin tracing library targeting each of 27 TADs of chromosome 22 (Chr22) was previously described (Wang et al., 2016), and this library was amplified using the same method as for the interphase DNA FISH probes (Liu et al., 2021). Primary hybridization of Chr22 probes was performed as described in the “Interphase DNA FISH” section. Sequential imaging was carried out using the custom microscope and microfluidics setup previously described (Liu et al., 2021). With little exception, imaging was performed as previously reported (Wang, 2016; Liu et al., 2021). To maximize the accuracy of TAD localization during data analysis, all images were acquired as multi-color Z-stacks, including consecutive imaging of Alexa Fluor 647 dyes (first 14 TADs of Chr22), ATTO 565 dyes (last 13 TADs of Chr22), and yellow-green fiducial beads for sequential image alignment at each Z-step. Images were collected with 0.5-sec per color per Z-height, 200-nm Z-step size, and 50 total Z-steps (10 µm total image depth). FISH signals were removed between rounds of sequential hybridization and imaging by 50 seconds of simultaneous illumination in the 647-nm and 560-nm color channels, as previously detailed^37,41^. Image analysis and chromatin trace assembly were performed using the MinaAnalyst package, available at https://campuspress.yale.edu/wanglab/mina-analyst/.

## Analysis of chromosome compaction

We calculated the inter-TAD distances between all pairs of TADs on chr22 for each cell type, and generated mean inter-TAD distance matrix by calculating the mean distance between each pair of TADs. To quantify the decompaction score, we then performed two-sided Wilcoxon signed rank test between the mean inter-TAD distances between untreated and osimertinib-treated cells and calculated FDR and fold change values, as previously described^45^.

## Determination of A and B compartment identity of TADs

We assigned A and B compartment identity to TADs using a previously published algorithm^37^. Briefly, we fitted a power-law function to the data points of the mean inter-TAD spatial distances versus their genomic distances across chromosome 22, which yielded the expected inter-TAD spatial distance for each pair of TADs according to their genomic distance. We then normalized the observed mean inter-TAD spatial distance by the expected spatial distance, yielding a normalized inter-TAD distance matrix. We then calculated the Pearson correlation coefficient between each pair of rows/columns in this matrix, generating a Pearson correlation matrix. We next applied principal component analysis to the Pearson correlation matrix, and took the coefficients of the first principal component as compartment scores. The compartment score profiles were validated with published ChIP-seq profiles in PC9 cells such that the compartment A regions are enriched in active histone modifications, gene density, and DNaseI accessibility. The following published ChIP-seq datasets from the ENCODE Project were used: ENCFF364RGE, ENCFF406PHZ, ENCFF605RZQ

## Analysis of long-range contact in A and B compartments

A long-range contact is defined when two non-adjacent TADs were less than 500nm apart from each other. We quantified the number of long-range contact events across all inter-TAD pairs and used the A and B compartments identifies to further quantify the number of AA, AB and BB long-range TAD contact events. Wilcoxon rank sum test was used to calculate statistical significance.

## RPKM and Peak Density Analysis

BAM files were converted to bed files using bamtobed from bedtools. Bedtools coverage was run with -a set to the BED file containing TAD coordinates and -b set to the BED file of the sequencing sample. METTL7A peak density was calculated by identifying peaks on chromosome 22 that belong to a particular TAD ID, summing the signalValue of peaks within such TAD ID, and normalizing the sum to the length of the TAD ID. RPKM and peak density were visualized in Matlab R2022b.

## Immunofluorescence

Cells were fixed with 4% paraformaldehyde and washed 3x with PBST (Filtered PBS + 0.1% Tween-20) and permeabilized for 20 min with PBS-Tx (PBS + 0.2% Triton X-100). Blocking was performed for 1 hour in PBST + 2% BSA. Samples were incubated with primary antibody overnight at 4°C in blocking buffer using the following antibodies: anti-MYC-tag 1:1000 (CST 2276S). Cells were washed 3x with PBST and incubated with secondary antibody for 1 hour at room temperature using the following secondary antibodies: Goat anti-Mouse IgG Alexa Fluor™ 488 (Thermo Fisher, A-11001) or Goat anti-Mouse IgG Alexa Fluor™ 568 (Thermo Fisher, A- 11031). Cells were washed with PBST and mounted in mounting media containing DAPI.

## Proximity ligation assay

Cells were fixed in 4% paraformaldehyde, washed 3x in PBS, and permeabilized for 20 min with PBS-Tx (PBS + 0.2% Triton X-100). Blocking was performed for 1 hour in PBST + 2% BSA. Samples were incubated with primary antibody overnight at 4°C in blocking buffer using the following antibodies: anti-MYC-tag 1:1000 (CST 2276S), PDS5A 1:500 (Thermo Fisher PA5- 57755), Rad21 1:1000 (Active Motif 91245). Cells were washed 3x with PBST. Proximity ligation assay (PLA) was performed using the Duolink® In Situ Red Starter Kit Mouse/Rabbit (Millipore DUO92101-1KT) according to the manufacturer’s specifications.

## RT-qPCR

RNA was extracted using the RNeasy Mini Kit (Qiagen, 74104O) and concentrations were measured via Nanodrop. One ug of RNA was reverse transcribed into cDNA using the iScript cDNA Synthesis Kit (BioRad 1708890) according to the manufacturer’s instructions. Each qPCR consisted of 5 ng cDNA, 1X iQ SYBR Green Supermix (BioRad 1708880), and 300 nM forward and reverse primers. Relative expression was calculated by the ddCt method relative to actin using the average ddCt values from 3 technical replicates. Each experiment was performed in biological triplicate. Primer sequences are listed in **Supplementary Table 4.**

## Immunoblotting

Approximately 2 x 10^6 cells were lysed with RIPA buffer (Thermo Fisher 89900) for 15 min on ice. Lysate was clarified by centrifugation at top speed for 30 min at 4°C. Protein concentration was quantified via BCA assay (Pierce). Approximately 20 ug of protein was mixed with 1X Lamelli buffer (BioRad 1610747) and 10% BME, boiled for 5 min, and loaded onto a 4-20% polyacrylamide gel (BioRad). Electrophoresis was performed at 100V for approximately 90 min. Gels were then transferred onto a PVDF membrane at 90V for 90 min in transfer buffer. Blots were blocked for 1 hour at room temperature in blocking buffer (5% milk in TBST [1X TBS, 0.5% Tween-20]) and incubated overnight at 4C in primary antibody diluted in blocking buffer. The following primary antibodies were used: anti-FLAG M2 1:1000 (Sigma F1804), anti-FLAG 1:1000 (Sigma F7425), anti-MYC-tag 1:1000 (CST 2276S), Histone H3 (CST 4499), PDS5A (Thermo Fisher, PA5-57755), Lamin B1 1:1000 (Proteintech 66095-1), Calnexin 1:1000 (CST 2679), alpha-Tubulin (CST 2144S), CoxIV (abcam ab16056), cRAF 1:1000 (CST 53745), and MET 1:1000 (CST 4560). Blots were washed 3X 5 min in TBST and incubated with secondary antibody (CST 7074P2 or 7076P2) diluted in blocking buffer for 1 hour at room temperature. Blots were washed 3X 5 min with TBST and visualized with SuperSignal Western Femto (Thermo 34096).

## Protein expression and purification

DNA encoding human METTL7a was inserted into a modified vector preceded by an N-terminal hexahistidine(His_6_)-MBP tag and a TEV cleavage site. BL21(DE3) RIL cells expressing the plasmids were induced by addition of 0.13 mM isopropyl β-D-1-thiogalactopyranoside (IPTG) when the cell density reached A_600_ of 1.1 and were grown at 16 °C overnight. The cells were harvested and lysed in buffer containing 50 mM Tris-HCl (pH 8.0), 1 M NaCl, 25 mM Imidazole, 10% glycerol and 1 mM PMSF. Subsequently, the fusion proteins were purified through a nickel column and size-exclusion chromatography. The purified protein samples were concentrated in 20 mM Tris-HCl (pH 7.5), 100 mM NaCl, 5% glycerol and 5 mM DTT, and stored at -80°C.

## Electrophoretic mobility shift assay

0.04 µM 60 nt DNA (ATTCTTCCGAGTTTTTTCCTCATCTCACTTCCAATACAGAAAGCTTGCCCCGCCCTTCCT and duplex (upper strand: ATTCTTCCGAGTTTTTTCCTCATCTCACTTCCAATACAGAAAGCTTGCCCCGCCCTTCCT; lower strand: AGGAAGGGCGGGGCAAGCTTTCTGTATTGGAAGTGAGATGAGGAAAAAACTCGGAAGAAT) was titrated with MBP-tagged METTL7A ranging from 0 to 0.8 µM in the 10-uL reaction mixture with the binding buffer containing 20 mM Tris-HCl (pH 7.5), 50mM NaCl, 5% glycerol and 5 mM DTT. Samples were incubated on ice for 20 min and then resolved on a 5% wt/v polyacrylamide gel (59:1 acrylamide: bis(acrylamide)), which was run at 100 V using 0.2x TBE (pH 8.3) running buffer for 50 min at 4°C. The gel was stained with SYBR™ Gold (Thermo) and visualized by ChemiDoc Imaging System (BioRad). The bound ratio (Fraction bound= bound/(bound+unbound)) from the band intensities for the free and protein-bound forms was quantified by using ImageJ. The dissociation constant (K_d_) was obtained by plotting the bound ratio versus protein concentration and fitting the curve to the Hill equation (Fraction bound =[protein]^h^/((K_d_)^h^+[protein]^h^) using GraphPad Prism 10.

## Acknowledgments

We thank C. Calderwood at the Yale Functional Genomics Core for help producing shRNA lentiviruses. We are grateful to P. Mischel, who generously provided COLO320-HSR and COLO320-DM cells for use in this study. We thank B. Bernstein for depositing PC9 CTCF, H3K9me2, and H3K9me3 ChIP-seq datasets to ENCODE, which were used in this study, and to E. Aiden for depositing PC9 Hi-C data to ENCODE. We thank T. Ardito and J. Wolenski for imaging support, R. Elmeskini and Z. Weaver Ohler from NCI Center of Advanced Preclinical Research (CAPR) for PDXs, N. Neuenkirchen for biochemistry support, M. Liu for chromosome tracing analyses, members of the Xiao lab for feedback, and the Yale Center for Research Computing for computational support. R.M.S. was supported by the NSF GRFP and 5T32HD007149-43. This work was supported by the Yale Lung SPORE.

## Author contributions

R.M.S. and A.Z.X. conceptualized and designed experiments. R.M.S. and E.G.S. performed imaging, sequencing, and cell proliferation/viability experiments and analyses. R.M.S., R.G., J.L., and N.K. performed biochemical experiments. E.G.S., J.R., and T.J. designed FISH and chromatin tracing probes. J.R. and R.M.S. performed chromatin tracing experiments. N.S. produced METTL7A knockout cells. N.R. generated PDXs used in Fig. 1 and Extended Data Figs 2-3. B.H. and M.A.M. generated PDXs and PDX RNA-seq data used in Fig. 2. J.S., K.P., S.W., and A.Z.X supervised the study. R.M.S., E.G.S., and A.Z.X wrote the paper with insights from all co-authors.

## Data Availability Statement

WGS has been deposited under PRJNA1065471 and will be released upon publication. RNA-Seq, ChIP-Seq, and CUT&RUN have been deposited in GEO database under accession number GSE254521.

## Captions for Supplementary Videos

**Supplementary Video 1.** 3D rendering of PC9 M7A-MYC expressing cells. Green labels M7A- MYC-tag and magenta labels nuclei.

**Supplementary Video 2.** 3D rendering of PC9-OR M7A-MYC expressing cells. Green labels M7A-MYC-tag and magenta labels nuclei.

**Supplementary Video 3.** 3D rendering of HCC827 M7A-MYC expressing cells. Green labels M7A-MYC-tag and magenta labels nuclei.

**Supplementary Video 4.** 3D rendering of HCC827-OR M7A-MYC expressing cells. Green labels M7A-MYC-tag and magenta labels nuclei.

**Supplementary Video 5.** 3D rendering of PC9-OR M7A-MYC expressing cells. Transparency is increased over the course of the video showing intra-nuclei foci. Green labels M7A-MYC-tag and magenta labels nuclei.

## Captions for Supplementary Tables

**Supplementary Table 1.** DESeq2 normalized counts table for RNA-seq analysis of cell lines used in this study.

**Supplementary Table 2.** Cutoffs used to generate Extended Data Fig. 2.

**Supplementary Table 3.** Raw mass spectrometry spectral counts and SAINTExpress analysis.

**Supplementary Table 4.** Primer sequences used throughout the study.

**Extended Data Fig. 1.**
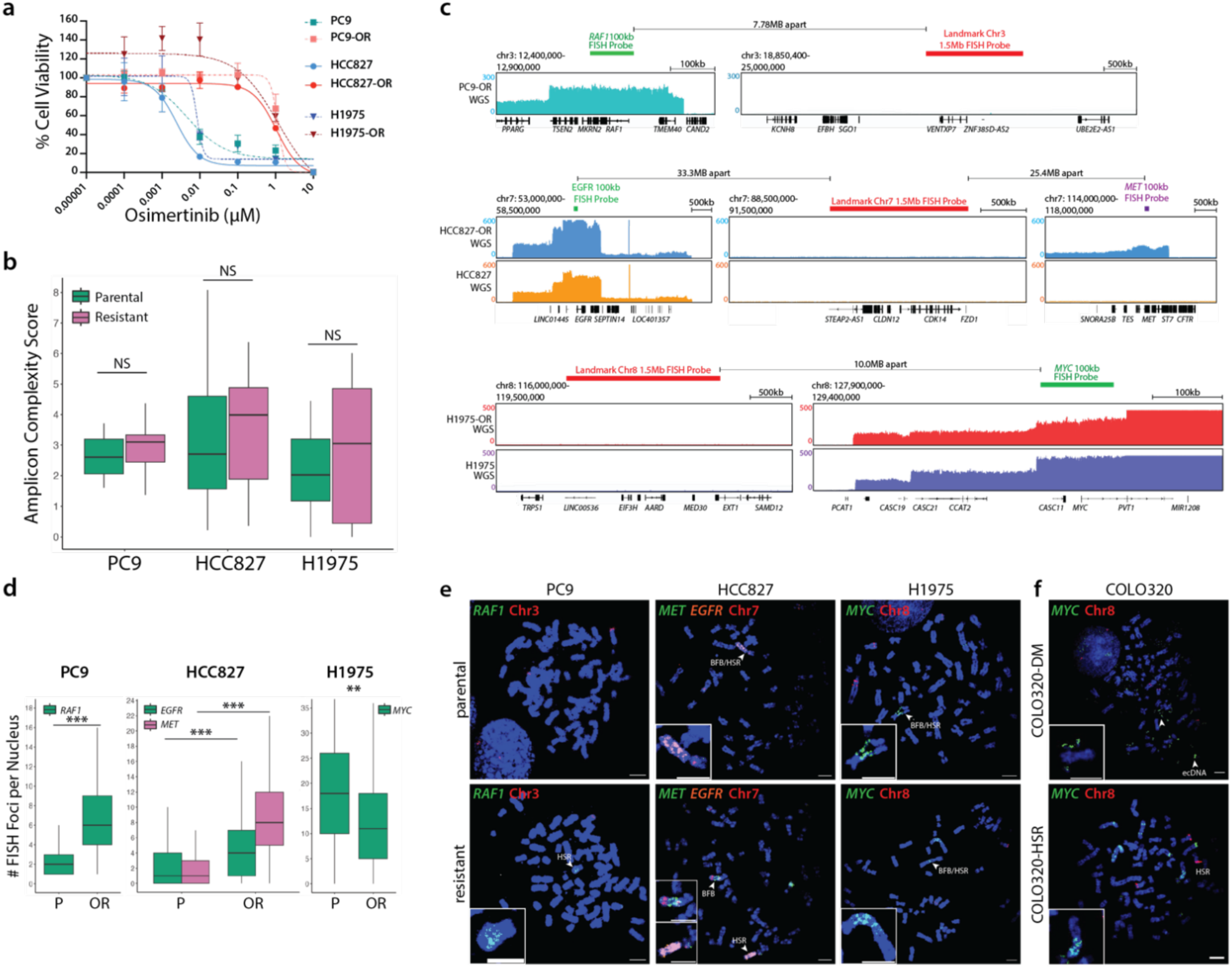
Oncogene amplification occurs in EGFR-mutant LUADs. **a**, Cell viability curves in EGFR-mutant LUAD cell lines PC9, HCC827, and H1975 comparing osimertinib sensitivity between parental cells and osimertinib-resistant (OR) cell lines. Error bars: standard deviation between three biological replicates. **b,** Amplicon complexity score increases, although not statistically significant (NS), in osimertinib resistant cells compared to parental cells. **c,** WGS IGV tracks with schematic of DNA FISH probe design strategy. DNA FISH probes were designed to target a 100 kb amplified locus or a 1.5 Mb unamplified control locus. **d,** Quantification of FISH foci from Fig. 1c. *P* values determined by two-sided Wilcoxon test. \*\**P* < 0.01, \*\*\**P* < 2.2 x 10^-16^. P: parental; OR: osimertinib-resistant. **e**, Representative metaphase FISH images in sensitive and resistant cells. Scale bars are 5 µm in all images except PC9-OR HSR inset, which has a 3 µm scale bar. **f**, Metaphase FISH images from COLO320-DM (top) and COLO320-HSR (bottom) which exhibit ecDNA- and HSR-like *MYC* amplicons, respectively.

**Extended Data Fig. 2.**
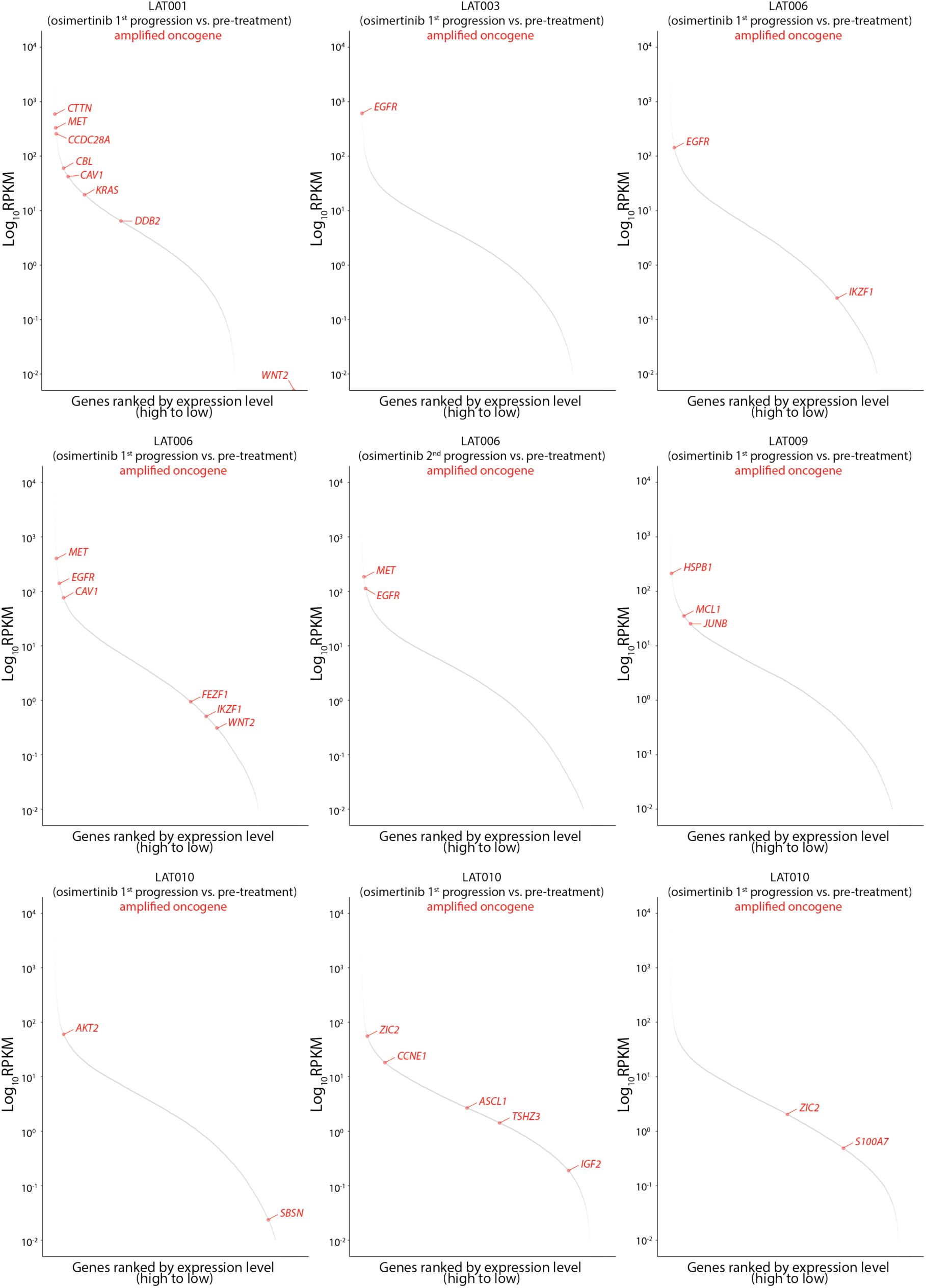
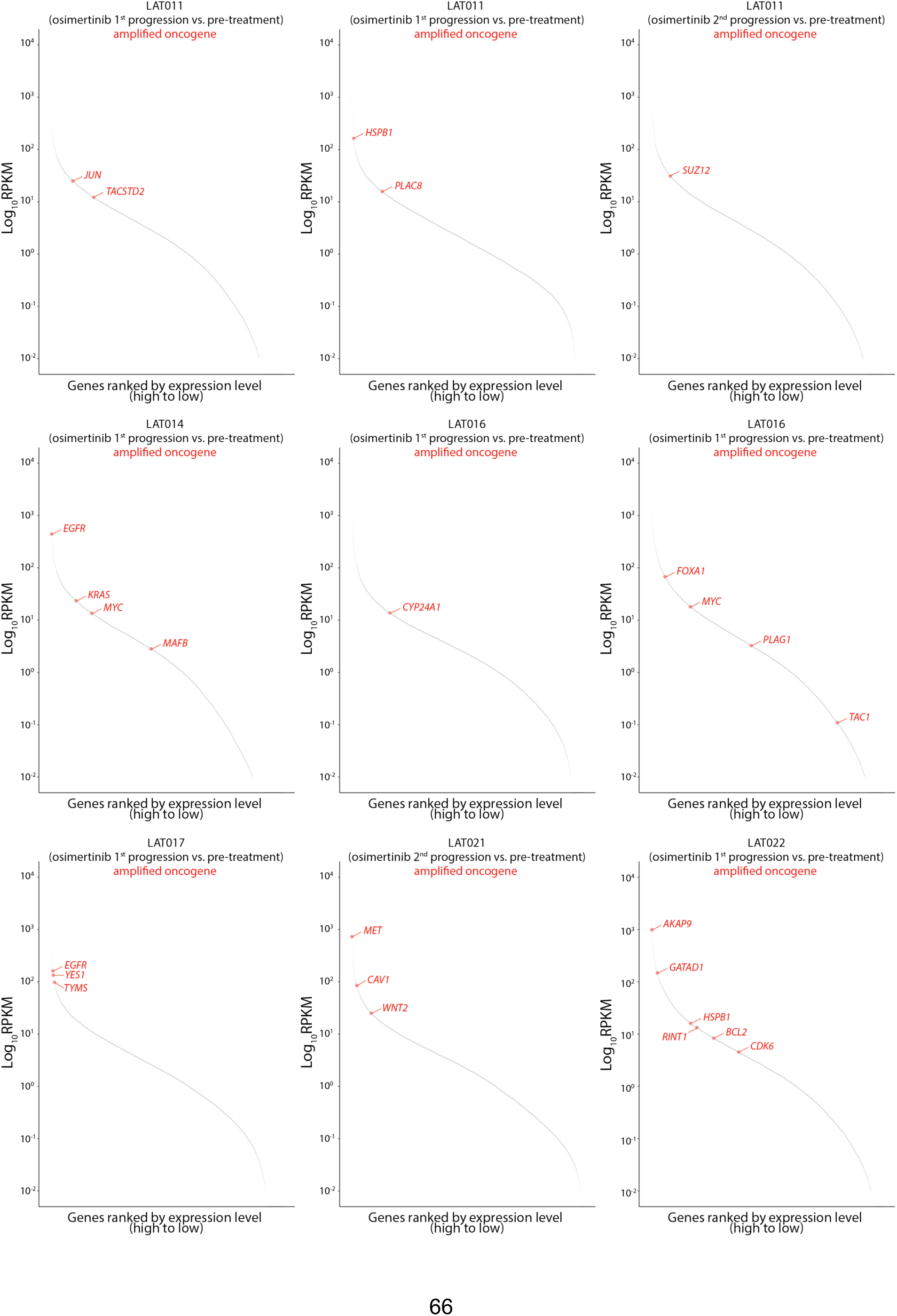
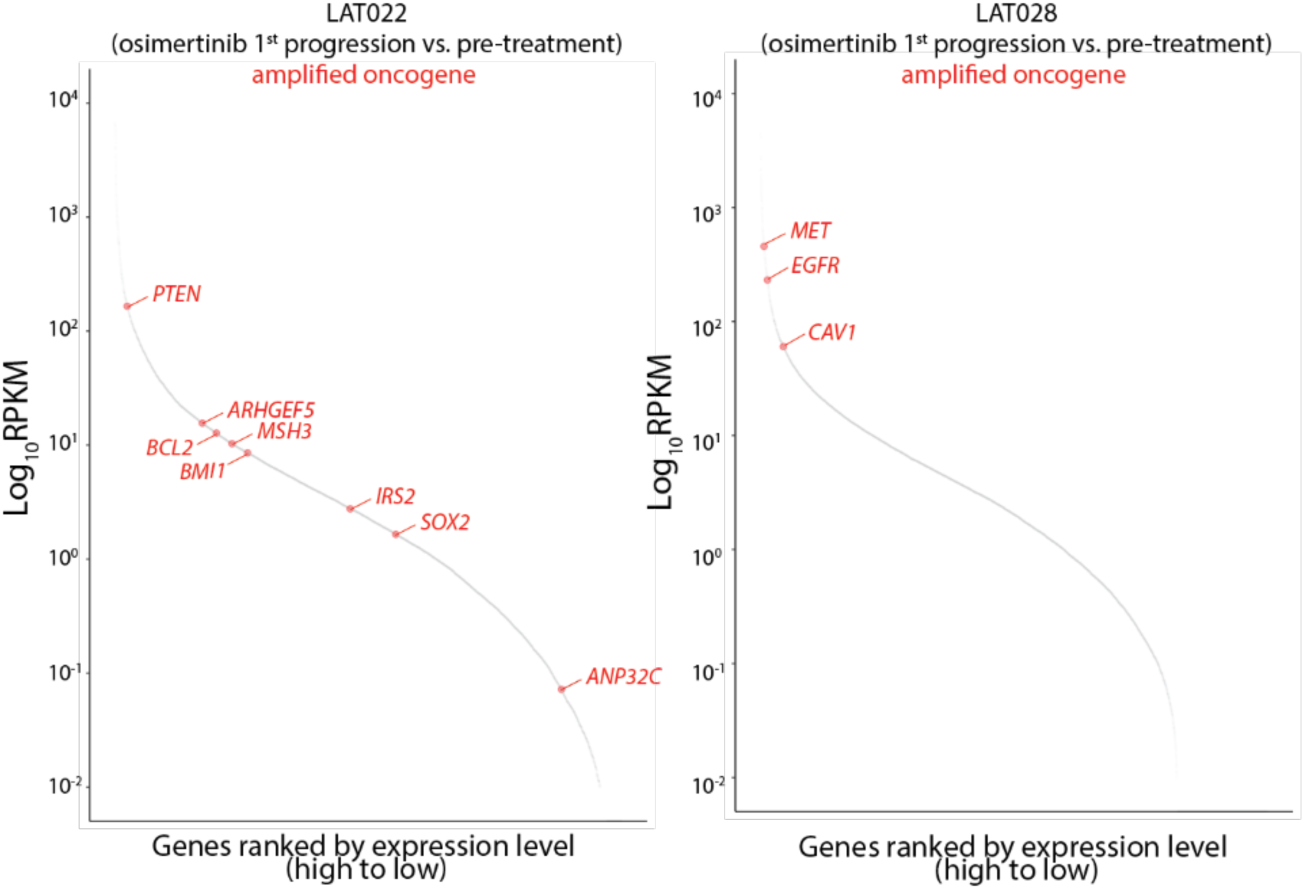
Oncogene amplification occurs in osimertinib-resistant EGFR-mutant LUADs from patients. Ranked RNA expression plots of osimertinib-treated tumors from NCT02759835. Copy number analysis was performed comparing 1^st^ or 2^nd^ progression tumors treated with osimertinib to untreated tumors using whole exome sequencing. Red dots are amplified oncogenes.

**Extended Data Fig. 3.**
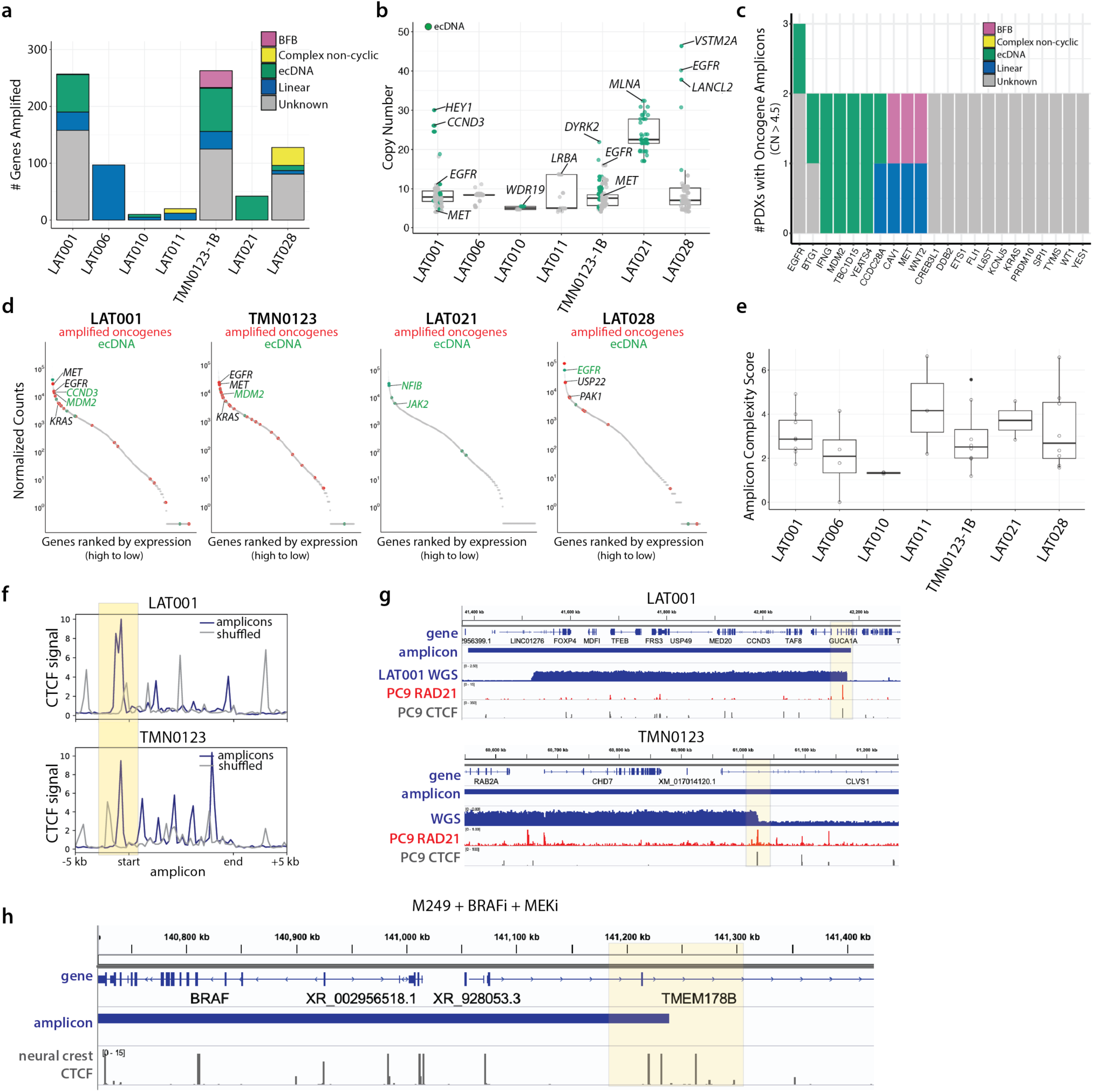
Oncogene amplicons have a pre-defined chromatin architecture. **a,** Breakdown of amplicon structures in PDX tumors derived from osimertinib-treated LUAD tumors. **b**, Copy number of amplified genes in PDX LUAD samples. Green: genes predicted to be amplified as ecDNA based on AA. **c,** Bar chart showing commonly amplified oncogenes across PDX tumor samples, colored by amplicon classification. **d**, Ranked RNA expression plots in osimertinib-treated PDX LUADs. Red dots: amplified oncogenes; green dots: ecDNA based on WGS/AA analysis. **e,** Amplicon complexity scores in PDX LUAD samples. **f,** Analysis of CTCF signal in PC9 cells based on the boundaries of amplicons from PDX tumors. CTCF is enriched at the boundaries of amplicons from PDX samples LAT001 and TMN0123. **g,** Example IGV sequencing tracks of PC9 CTCF signal (grey) and PC9 RAD21 signal (red) correlated with the boundaries of the amplicons from PDX tumors LAT001 (top) and TMN0123 (bottom). Both CTCF and RAD21 are enriched at one amplicon boundary. **h,** Example IGV sequencing track of CTCF signal from neural crest cells correlated with the boundary of the BRAF amplicon in M249 cells resistant to BRAFi and MEKi.

**Extended Data Fig. 4.**
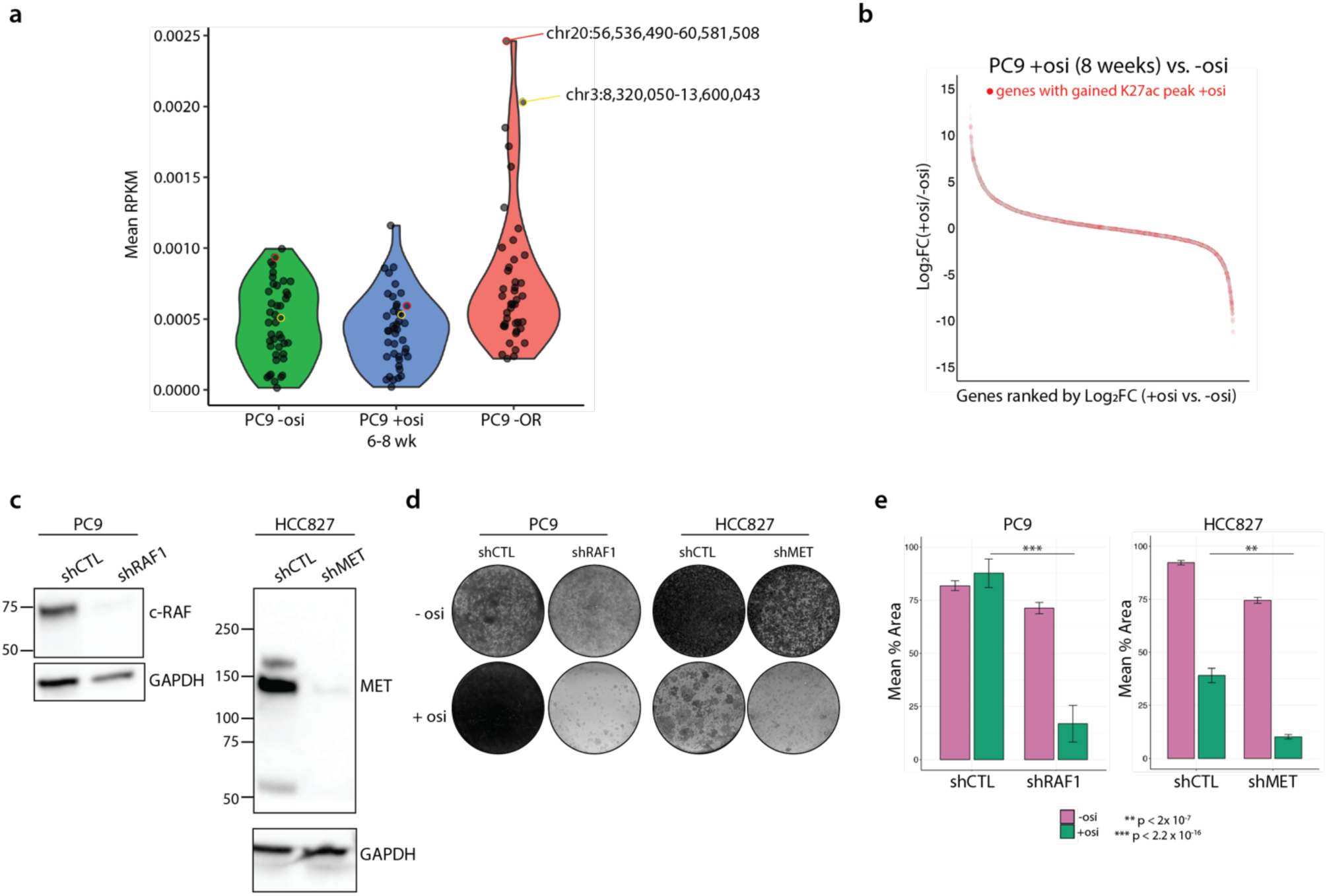
Chromatin priming precedes oncogene amplification. **a,** Genome coverage (RPKM) based on H2A CUT&RUN in PC9 cells treated without osimertinib or with osimertinib for 6-8 or 12+ (PC9-OR) weeks. Each dot represents an amplicon locus determined by AA in PC9-OR cells. Signal is normalized for sequencing depth. **b,** Ranked RNA expression plot showing Log_2_FC between PC9 cells treated with osimertinib for 8 weeks compared to parental cells. Genes that have gained an H3K27ac peak upon osimertinib treatment are highlighted in red. **c,** Western blots showing shRNA-mediated knockdown efficiency of RAF1 in PC9 cells (left) and MET in HCC827 cells (right). **d,** Colony formation assay in PC9 (left) and HCC827 (right) cells expressing shRNAs targeting RAF1 or MET, respectively, or a non-targeting control. Cells were treated with osimertinib for approximately 5 weeks and fixed and stained with crystal violet. **e,** Quantification of (d). P-values determined by an unpaired t-test. Error bars represent standard deviation from three biological replicates.

**Extended Data Fig. 5.**
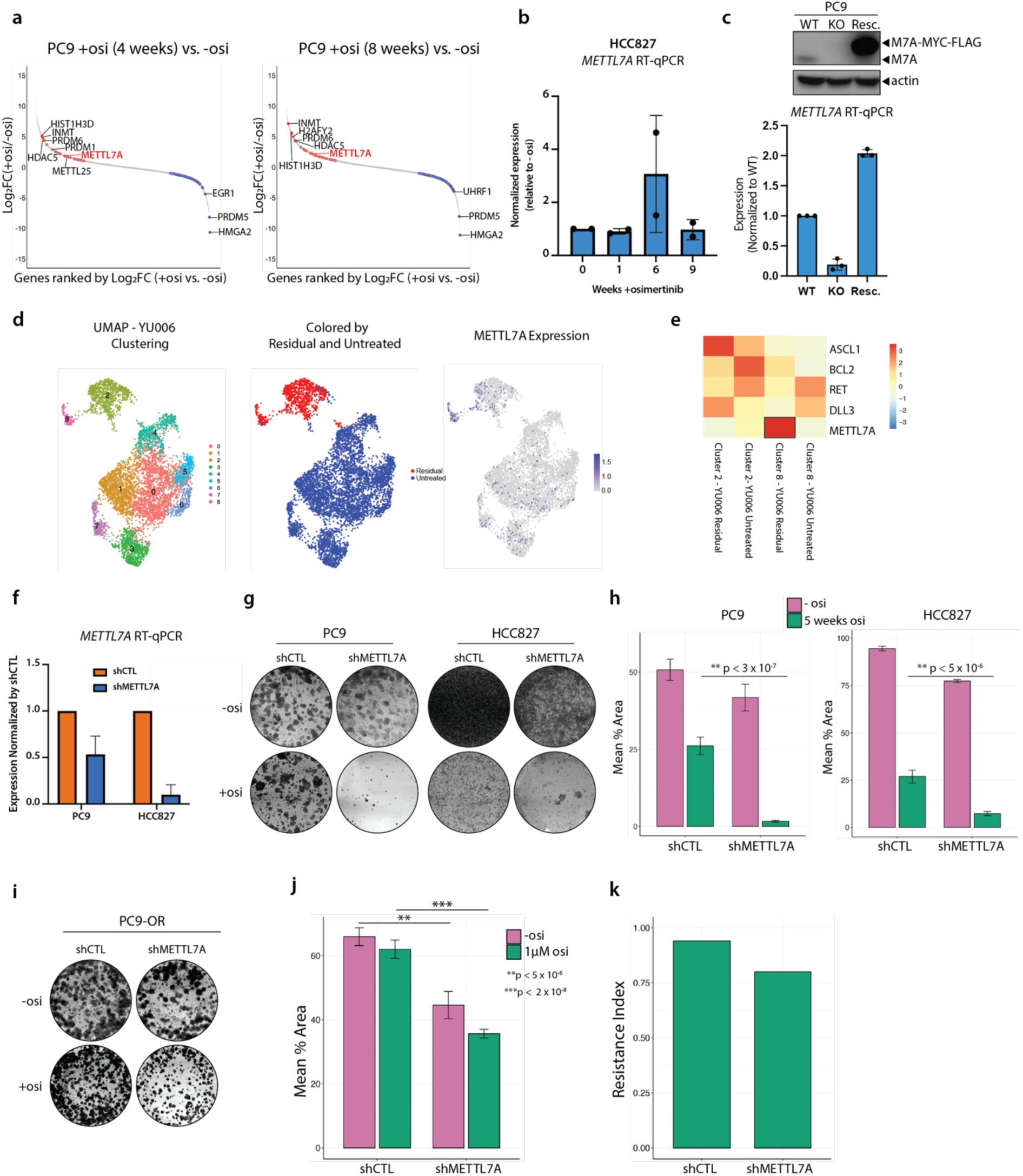
**METTL7A promotes the acquisition of osimertinib-resistant LUAD**. **a,** Ranked RNA expression plot showing the differentially expressed (P_adj_ < 0.001) putative and known epigenetic factors that are upregulated (red, log_2_FC > 1) or downregulated (blue, log_2_FC < 1) after 4 (left) and 8 (right) weeks of osimertinib treatment. **b,** RT-qPCR of *METTL7A* in HCC827 cells treated with osimertinib at the indicated time points from two biological replicates. **c,** Top: Western blot in PC9 *METTL7A* WT, KO, or KO cells rescued by overexpressing METTL7A-MYC-FLAG. Bottom: RT-qPCR of *METTL7A* in *METTL7A* WT, KO, or KO cells rescued by overexpressing METTL7A-MYC-FLAG. **d,** (left) UMAP of YU-006 PDX samples colored based on clusters, (middle) colored by condition, (right) colored by *METTL7A* expression. **e,** Heatmap visualization showing *METTL7A* expression in clusters 2 and 8 from (d) along with previously identified factors such as ASCL1, that pre-exist in YU-006 untreated tumor. **f,** RT-qPCR of *METTL7A* in PC9 and HCC827 cells expressing a nontargeting control shRNA (shCTL) or an shRNA targeting *METTL7A.* **g,** Colony formation assay in PC9 and HCC827 cells expressing an shRNA targeting METTL7A or a non-targeting shCTL. shMETTL7A cells fail to form osimertinib-resistant colonies after approximately 4 weeks. **h**, Quantification of (g). Error bars represent standard deviation between biological triplicates. Significance determined by unpaired t-test. **i,** Colony formation assays in PC9-OR shCTL and shMETTL7A cells treated with or without osimertinib. **j,** Quantification of (i). **k,** Resistance index (mean percent area -osi/mean percent area +osi). Error bars represent standard deviation between biological triplicates with three technical replicates per biological replicate.

**Extended Data Fig. 6.**
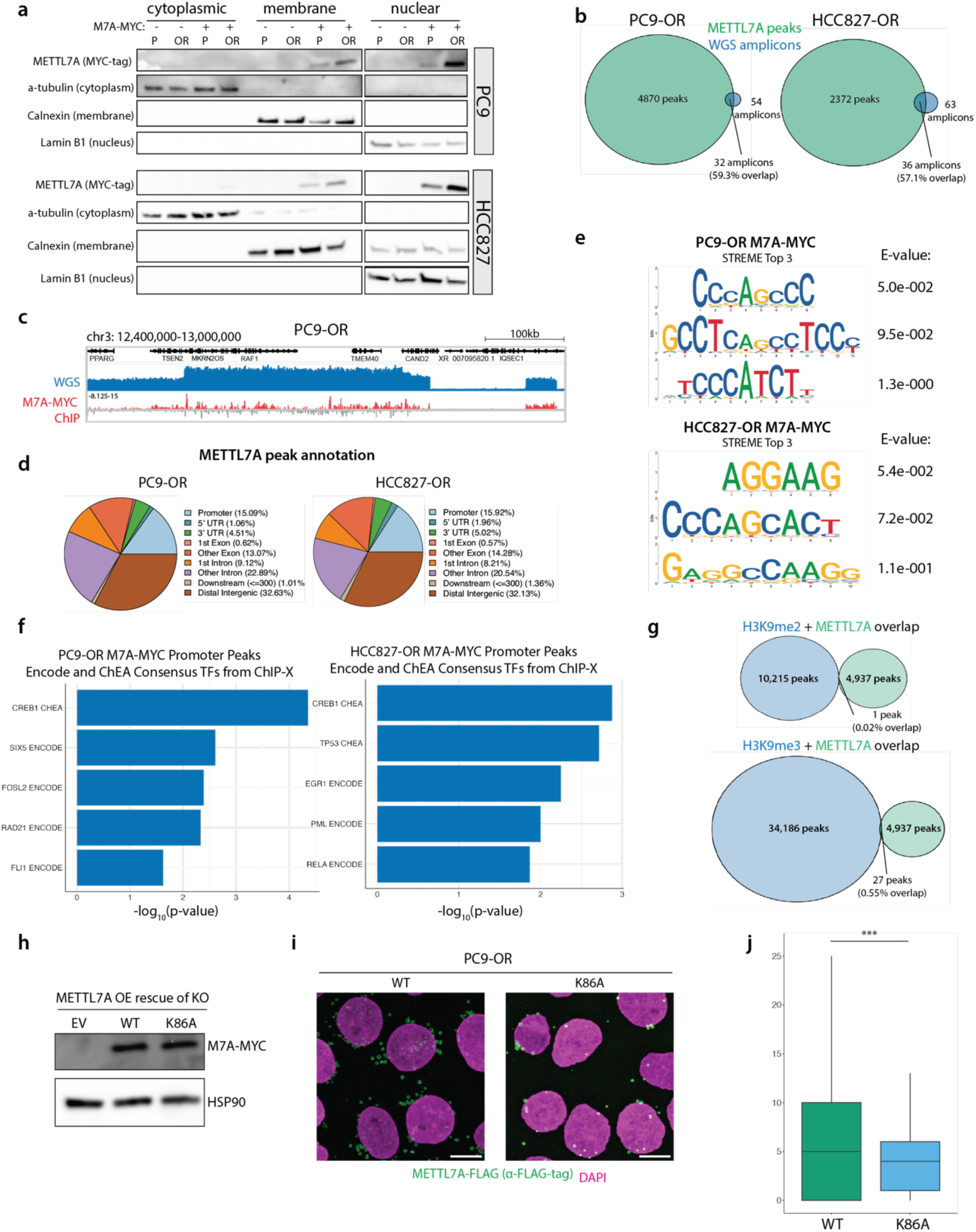
METTL7A binds to amplified oncogenes. **a,** Subcellular fractionation of PC9 and HCC827 cells that express MYC-tagged METTL7A. Wildtype cells that do not express METTL7A-MYC were used as negative controls. α-tubulin, Calnexin, and Lamin B1 antibodies were used to ensure the purity of each indicated fraction. P: parental, OR: osimertinib-resistant. **b,** Overlap between METTL7A ChIP peaks and amplicons determined from WGS. **c,** Example IGV track of METTL7A enrichment over amplicons in PC9-OR cells. BigWig signal is normalized by ChIP input. **d,** METTL7A peak annotation in PC9-OR and HCC827-OR cells. **e,** METTL7A motif analysis via STREME. **f,** GO analysis of METTL7A promoter peaks in PC9-OR and HCC827-OR cells. **g,** Venn diagrams showing overlap between METTL7A and H3K9me2 (top) and H3K9me3 (bottom). **h,** Western blot of rescue constructs overexpressed in the METTL7A KO background. EV = empty vector. **i,** Representative immunofluorescence images of METTL7A-FLAG tag in PC9-OR cells expressing either WT METTL7A-FLAG-tag or K86A-mutant METTL7A-FLAG-tag. Scale bar is 10 microns. **j,** Quantification of nuclear METTL7A-FLAG-tag foci. ***p < 0.001.

**Extended Data Fig. 7.**
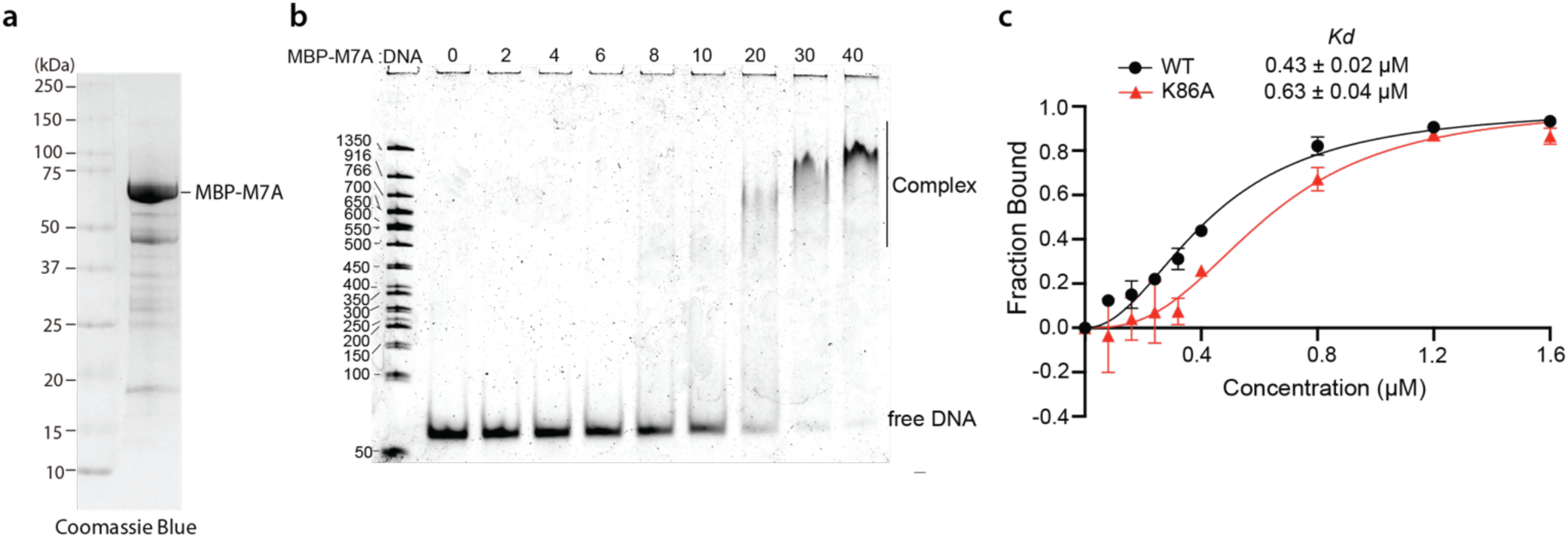
Recombinant METTL7A binds to DNA *in vitro*. **a,** Coomassie gel showing MBP-tagged METTL7A purified from *E. coli.* **b,** Gel shift assay shows MBP-M7A- dsDNA complex formation at increasing ratios of MBP-M7A-to-dsDNA. **c,** Quantification of gel shift assays with recombinant MBP-M7A WT and MBP-M7A-K86A. The Kd value was calculated based on the fraction of bound dsDNA at increasing concentrations of MBP-METTL7A.

**Extended Data Fig. 8.**
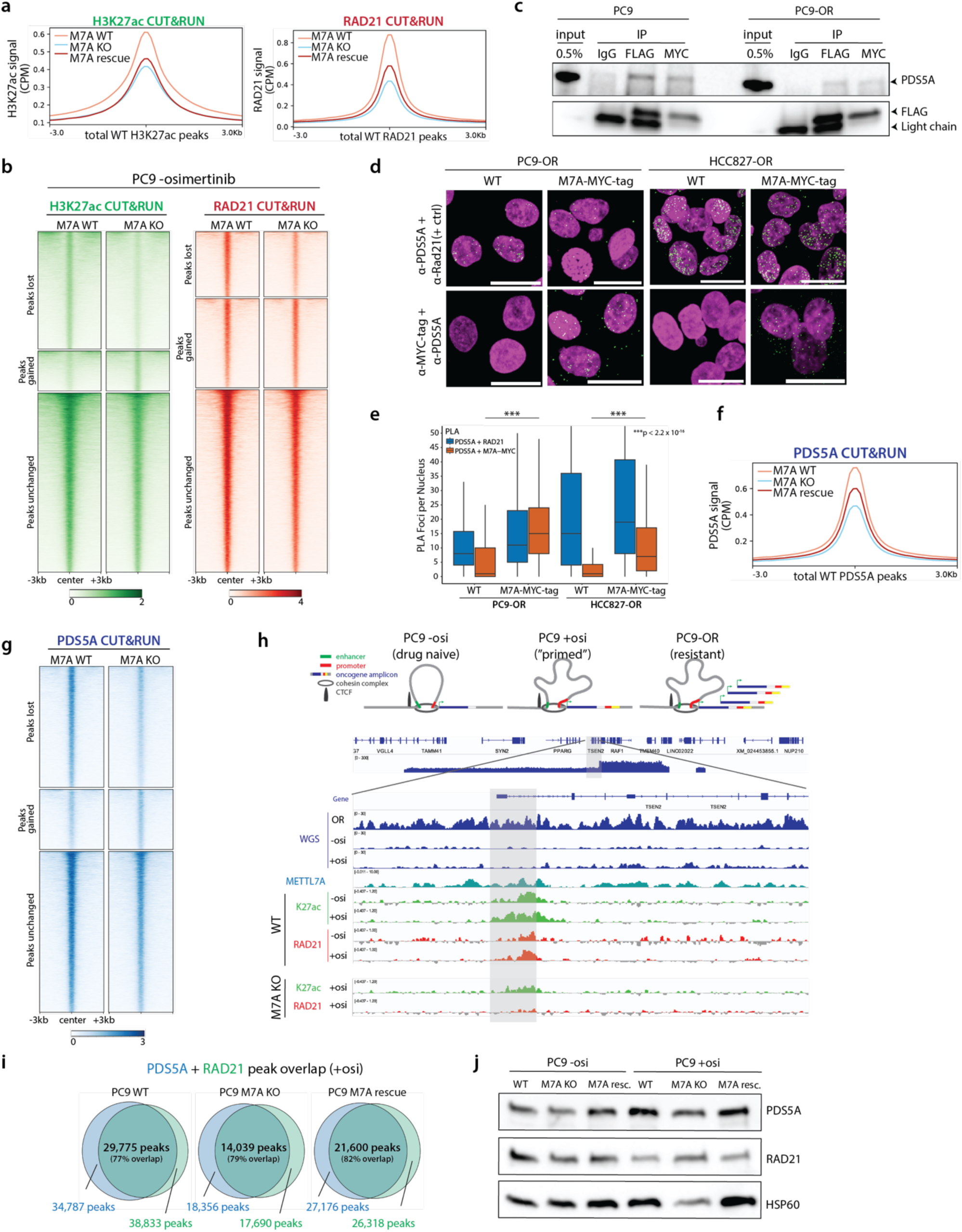
METTL7A affects the deposition of cohesin components. **a,** Metaplots of H3K27ac and RAD21 CUT&RUN in PC9 cells treated with osimertinib in the METTL7A (M7A) WT, KO, or rescue background. **b,** Heatmaps of H3K27ac and RAD21 CUT&RUN data in PC9 cells treated without osimertinib (parental cells). Signal is centered on peaks that were lost, gained, or unchanged upon M7A KO. CUT&RUN signal is normalized by CPM, H2A signal is subtracted, and mean bigWig signal between two biological replicates is shown. **c,** METTL7A-MYC-FLAG immunoprecipitation using antibodies against IgG, MYC, or FLAG followed by PDS5A western blotting shows binding between METTL7A-MYC-FLAG and PDS5A relative to 0.5% input. Anti-FLAG blotting shows the efficiency of each IP. **d,** Proximity ligation assays in PC9-OR and HCC827-OR cells show that METTL7A and PDS5A are in close proximity. PDS5A and RAD21 antibodies were used as a positive control. Wildtype (WT) cells that do not express the METTL7A-MYC-FLAG fusion protein were used as a negative control. Green signal: PLA foci; magenta: DAPI. Scale bar: 20 µm. **e,** Quantification of 7d. \*\*\**P* < 2.2 x 10^-16^**. f,** Metaplot of PDS5A CUT&RUN in PC9 cells treated with osimertinib in the METTL7A (M7A) WT, KO, or rescue background. **g,** Heatmaps of PDS5A CUT&RUN data in PC9 cells treated without osimertinib (parental cells). Signal is centered on peaks that were lost, gained, or unchanged upon M7A KO. CUT&RUN signal is normalized by CPM, H2A signal is subtracted, and mean bigWig signal between two biological replicates is shown. **h,** (top) Schematic depicting how the chromatin structure is “primed” during the acquisition of resistance prior to the development of resistant cells with oncogene amplification. (bottom) IGV tracks from WGS (dark blue), METTL7A ChIP in PC9-OR METTL7A-MYC cells (teal), H3K27ac CUT&RUN (green), and RAD21 CUT&RUN (red). H3K27ac and RAD21 are gained at the boundaries of the “future amplicons” (loci that are amplified in PC9-OR cells but not in “primed” cells). METTL7A KO leads to reduced H3K27ac and RAD21 at these loci. **i,** Overlap of PDS5A and RAD21 CUT&RUN peaks in cells treated with osimertinib. Percent overlap indicates percentage of overlapped peaks compared to total number of RAD21 peaks. **j**, Western blots of PDS5A and RAD21 show that changes in PDS5A and RAD21 deposition are not due to changes in protein levels.

**Extended Data Fig. 9.**
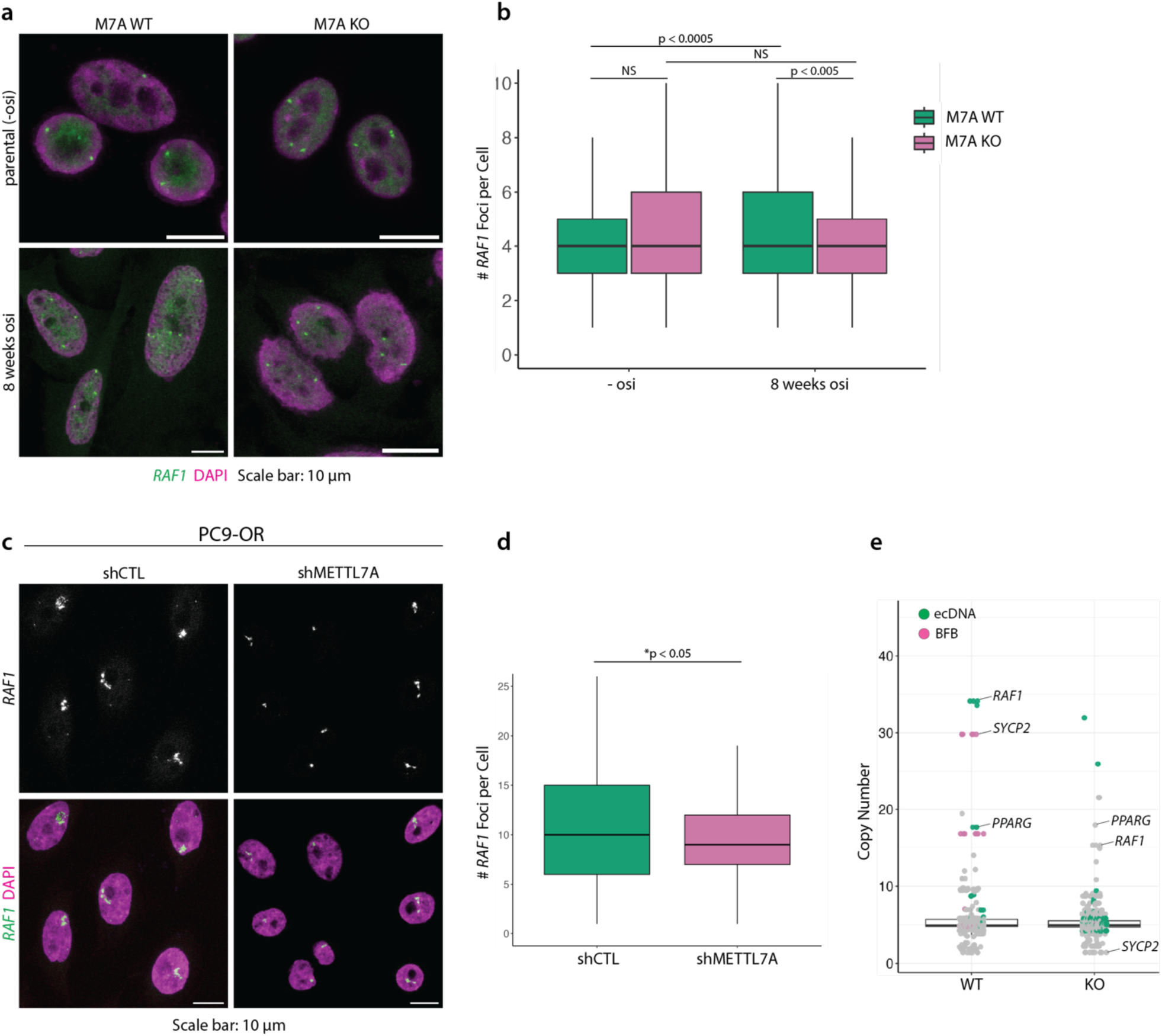
METTL7A affects gene copy number. **a,** *RAF1* FISH in PC9 WT and *METTL7A* KO parental cells and cells treated with osimertinib for 8 weeks. **b,** Quantification of RAF1 DNA FISH in **(a). c,** *RAF1* FISH in PC9-OR shCTL and shMETTL7A cells. **d,** Quantification of RAF1 DNA FISH in (c). **e,** WGS and AA analysis of PC9- OR WT and KO cells shows a decrease in average copy number upon *METTL7A* depletion.

**Extended Data Fig. 10.**
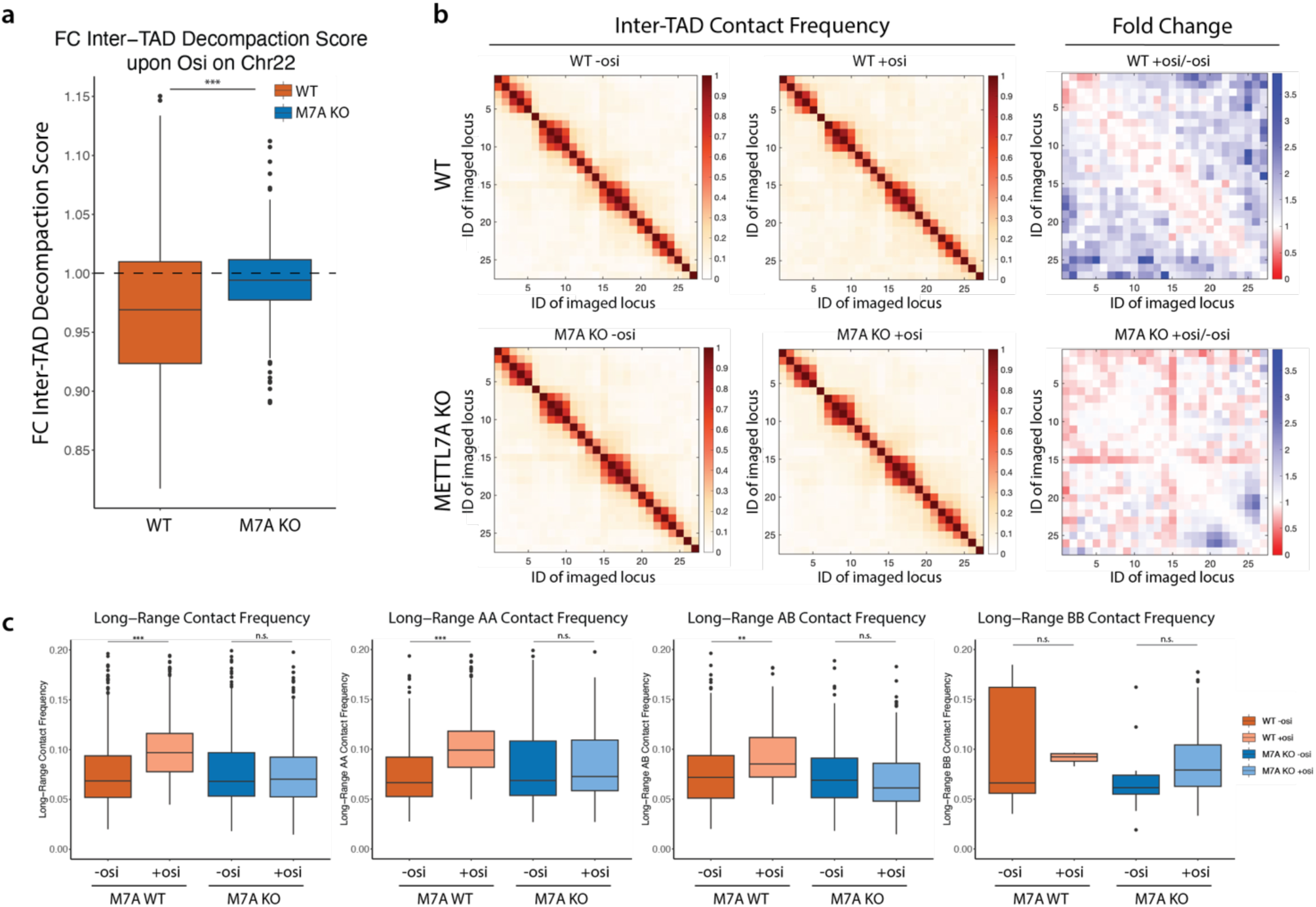
METTL7A affects chromatin compaction as cells acquire resistance to osimertinib. **a,** Fold change of inter-TAD decompaction score upon osimertinib (osi) treatment in PC9 WT and METTL7A (M7A) KO cells. **b,** Chr22 tracing reveals increased long-range inter-TAD contacts in wildtype cells treated with osimertinib compared to METTL7A KO cells. **c,** Quantification of long-range inter-TAD contact frequency across all long-range inter- TAD pairs, AA inter-TAD pairs, AB inter-TAD pairs, and BB inter-TAD pairs. (*** *P* < 0.001; ** P < 0.01; n.s. not significant).

